# Ipsilateral restriction of chromosome movement along a centrosome, and apical-basal axis during the cell cycle

**DOI:** 10.1101/2023.03.27.534352

**Authors:** Pingping Cai, Christian J. Casas, Gabriel Quintero Plancarte, Lisa L. Hua, Takashi Mikawa

## Abstract

Little is known about how distance between homologous chromosomes are controlled during the cell cycle. Here, we show that the distribution of centromere components display two discrete clusters placed to either side of the centrosome and apical/basal axis from prophase to G_1_ interphase. 4-Dimensional live cell imaging analysis of centromere and centrosome tracking reveals that centromeres oscillate largely within one cluster, but do not cross over to the other cluster. We propose a model of an axis-dependent ipsilateral restriction of chromosome oscillations throughout mitosis.

## Introduction

Chromosomes are highly organized structures within the nucleus of the cell [1]. Multiple studies have investigated the distribution of chromosomes across various human cell types, and have proposed different models for individual chromosome organization [2]. Interphase chromosome organization is radially distributed in the nucleus in what was originally thought to be random patterns [3, 4], but is increasingly correlated with factors such as chromosome size, gene density, or heterochromatic content [5–9]. Variations across different cell types [10], and nuclear morphologies [11] have also complicated the interpretation for one consistent model for chromosome organization in human cells. Although chromosomes 18 and 19 are similar in size, they differ in gene density [12, 5]. The study proposed that gene density determined chromosome positioning rather than size [12, 5]. Cell types with different nuclear morphologies and different fixation techniques have also contributed to conflicting studies [13–16]. In addition, the organization of homologous chromosomes in somatic cells were reported to be nonrandom [17]. However, it remains largely unknown how distance between the homologous chromosomes is regulated.

We have previously demonstrated the presence of a haploid (1n) chromosome set organization in primary human umbilical vein endothelial cells (HUVECs) [17]. We found that all homologous autosomal pairs, and the sex chromosomes were segregated, or antipaired, across the centrosome axis at metaphase/early anaphase in HUVECs [17]. This segregation pattern defined a haploid chromosome set (1n) per nuclear hemisphere organization, as defined by the centrosome axis, and was shown to be consistent across multiple cell types [17]. This study raises the question as to how the individual chromosomes within a haploid set remain together, and are excluded from the other haploid set.

Here, we report a novel distribution of centromeric DNA and protein components along the centrosome axis between chromosome sets in metaphase cells using fixed cell marker analysis. We use 4-Dimensional time lapse microscopy to reveal that centromere movements are largely restricted in an ipsilateral axis-dependent manner. Surprisingly, we find the axis-based restriction is present from mitosis onset to G_1_ interphase, even as the mitotic spindle is assembling, and disassembling. This suggests other non-spindle regulatory mechanisms that may maintain individual chromosome positions throughout the cell cycle. The ipsilateral-based restriction of chromosome organization and dynamics provides a framework to investigate mechanisms of haploid set organization.

## Results

### Distribution of centromeric DNA satellite sequences and the CENP-B protein in fixed primary human cells at metaphase

HUVECs were previously used to identify the haploid chromosome set-based, or antipairing, organization along the centrosome axis in metaphase/early anaphase cells (S1 Fig) [17]. HUVECs are adherent primary endothelial cells that demonstrate apical/basal polarity when grown in culture [18]. The polarized HUVECs allowed us to establish an axial coordinate system using the centrosomes as subcellular markers to define the x-axis; the optical path to define the z-axis; and the line perpendicular to both the x-, and z-axes to define the y-axis [17] (S2 Fig).

Human centromeres largely consist of α-satellites with cenpb DNA sequences, and this centromeric DNA has been implicated in interchromosomal linkages [19–30]. Early studies using electron microscopy, and Giemsa staining of metaphase spreads first described the existence of interchromosomal linkages between separate heterologous chromosomes in human, mouse, and Chinese Hamster cells [31–34]. These interchromosomal linkages were shown to stain positive with Hoechst DNA dye [35]. Subsequent micromanipulation experiments demonstrated that chromosomes moved in concert when pulled under tension, and could be mechanically dissected together as “beads on a string” in human cell lines [35–36]. Interchromosomal linkages were also corroborated by DNA Fluorescence *in situ* hybridization (FISH) assays in metaphase spread preparations of human and mouse cells [19–20], and have been shown to stain positive for sequences found within the peri/centromeric DNA regions of the chromosomes [19–30]. Past studies utilized metaphase spreads, which may damage the 3D endogenous spatial organization of chromosomes, and linkage structures [19–21]. Although the presence of these interchromosomal centromeric-based linkages have been reported, their function in the cell remains unknown.

To visualize the organization of centromeric components previously described for interchromosomal linkages, we performed DNA FISH in HUVECs, probing for DNA α-satellite and CENP-B box (cenpb) sequences (Fig 1A-1E). Immunofluorescence FISH (ImmunoFISH) assays for these sequences were conducted to locate centromeres in our fixed cell analysis. Visualization of the centrosomes by γ-tubulin immunofluorescence (IF) was also used to detect the mitotic stage of individual mitotic cells, which allowed us to map the haploid chromosome sets across the centrosome axis at metaphase and anaphase.

**Fig 1.**
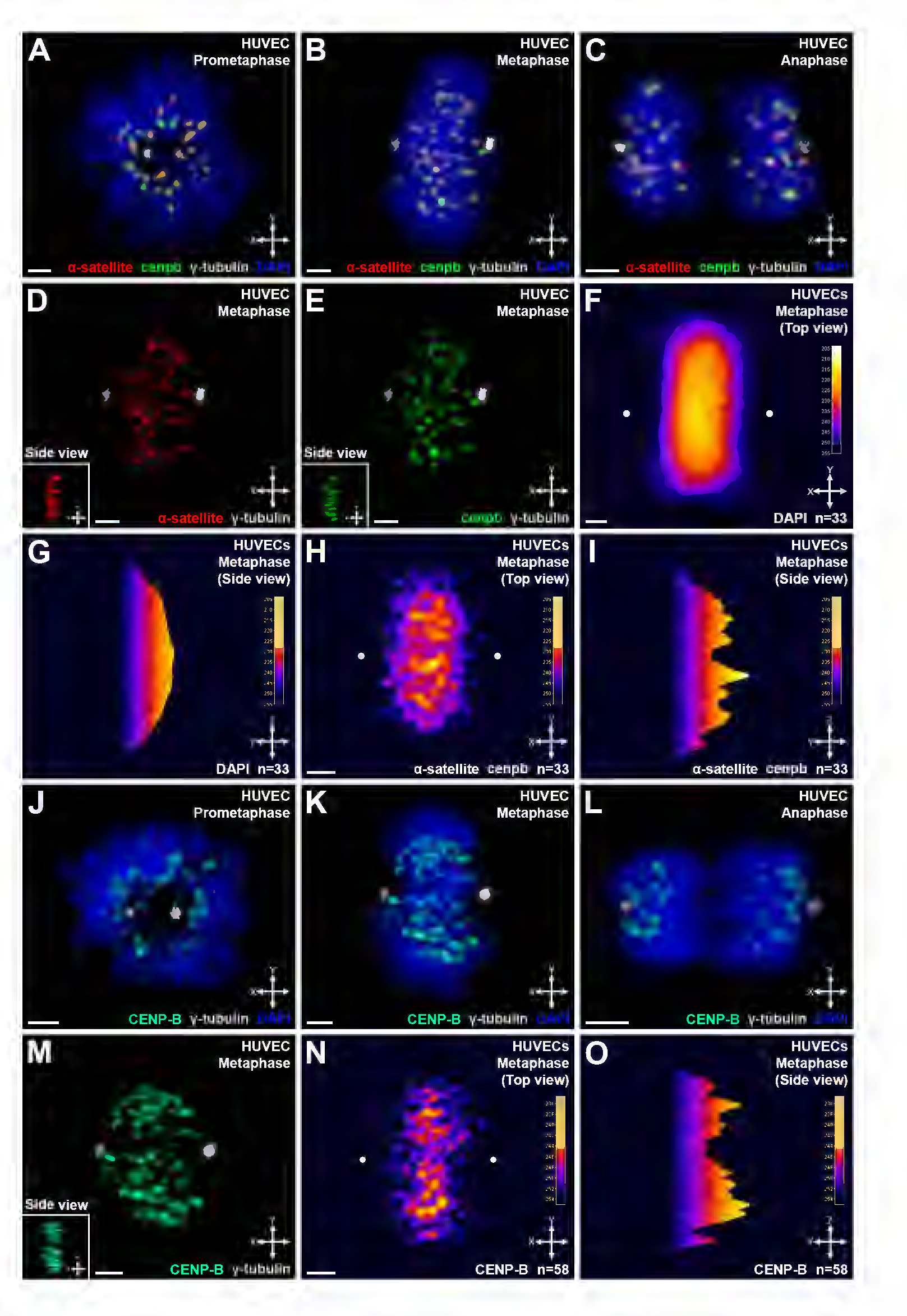
Distribution of centromeric DNA satellite sequences and the CENP-B protein in fixed primary human cells at metaphase. **(A)** A human umbilical vein endothelial cell (HUVEC) at prometaphase hybridized with DNA FISH for α-satellite (red), and cenpb (green), and stained with a γ-tubulin antibody (gray), and DNA counterstain (DAPI, blue). **(B, C)** As in (A), but at metaphase (B) and anaphase (C). **(D, E)** As in (B), but of the α-satellite (D), or cenpb (E) channel. (Inset) Side view. **(F)** Top view of a comprehensive heatmap for DAPI DNA stained HUVECs at metaphase. HUVECs were aligned along the centrosome axis (n=33 cells). White dots denote centrosome positions used for alignment. **(G)** As in (F), but side view. **(H, I)** As in (F, G), but of both α-satellite and cenpb DNA satellite staining (n=33 cells). **(J-L)** As in (A-C), but immunolabeled for CENP-B protein (green) and γ-tubulin (white). **(M)** As in (K), without DNA counterstain. (Inset) Side view. **(N,O)** As in (F,G), but for CENP-B protein staining (n=58 cells). Scale bars: 2 µm.

Throughout mitosis in intact human cells, DNA satellite sequences (α-satellite and cenpb) were readily identified (Fig 1A-C). Satellite staining was visible between multiple foci within the DNA mass. In contrast, interchromosomal linkages were difficult to be identified, except for metaphase at which a unique pattern of centromeric DNA components emerged due to lesser chromosome compaction as compared to prometaphase and anaphase (Fig 1D,E) [37–39]. Therefore, HUVECs at metaphase were examined for centromeric characterization and analysis. They frequently displayed continuous centromeric DNA satellite staining across the centrosome axis (Fig 1D,E, S1 Video). Individual channels for α-satellite (Fig 1D) and cenpb sequences (Fig 1E) revealed a region of low DNA satellite staining (n=23/33 cells). This region was on average 0.77 µm +/- 0.19 µm s.d. in width, and persisted across the metaphase DNA mass.

To test whether this unique pattern of centromeric DNA satellite staining is conserved across multiple HUVECs at metaphase, we constructed a comprehensive heatmap. The heatmap of DNA satellite staining was constructed by aligning individual cells along their centrosome axis using the axial coordinate system as defined by Hua and Mikawa [17] (Fig 1F-I). The heatmap provides objective quantification of the signal to noise intensity for the centromeric DNA satellite staining. Individual centromeres can oscillate at a range of ∼2 µm min^-1^ during metaphase in live cells [40]. Therefore, we applied a rotational analysis for each cell along different axes to determine if a consistent centromeric distribution pattern could be identified (S3 Fig). We found a consistent centromeric satellite staining pattern following an average of ∼18° rotation along the centrosome axis. In addition, some cells also exhibited multiple regions of low DNA satellite staining (n=30/65 cells) (S4 Fig). Our constructed DNA satellite heatmap of both the α-satellite and cenpb sequences showed a valley/depression coincident with the centrosome axis (Fig 1H,I). For comparison, a heatmap of DNA staining is shown (Fig 1F,G). These results support a clearly discrete distribution of centromeric linker components with two peaks and a valley of DNA satellites that is conserved among HUVECs along the centrosome axis.

In addition to DNA satellite sequences, the CENP-B protein is a centromeric component and directly interacts with the cenpb DNA satellite sequence [19]. To test whether the CENP-B protein showed a similar localization pattern as the satellite sequences, we next performed IF for CENP-B, and γ-tubulin in HUVECs (Fig 1J-M). Similar to our DNA satellites survey, two peaks of CENP-B positive centromeric staining were visible at metaphase (Fig 1N,O). A low region of CENP-B staining with an average width of 0.81 µm +/- 0.32 µm s.d. also persisted across the metaphase DNA mass in individual cells (Fig 1M, n=45/58 cells). The CENP-B heatmap also exhibited a valley/depression coincident with the centrosome axis (Fig 1N, O). These data support a conserved pattern for a centromeric protein component, CENP-B, along the centrosome axis in HUVECs. The set of data demonstrates a discrete distribution pattern of centromeric components with two sub-groups that flank a plane along the centrosome-centrosome, and apical/basal axes. This distinct pattern of centromeric components suggests the presence of a mechanism that restricts their random movement and/or distribution in the cell.

### Ipsilateral, and apical/basal axis-based restriction of centromere movements along the centrosome axis from metaphase to anaphase

The discrete distribution pattern of centromeric components across the centrosome axis in fixed cells suggested that centromeric movements/fluctuations may not be random, and be restricted instead. To test this possibility, we employed a 4D high resolution confocal live imaging analysis using the human Retinal Pigment Epithelial-1 (RPE1) cell line expressing CENP-A/centrin1-GFP, which labels the individual centromeres, and centrosomes, respectively [37]. This system allowed us to track centromere motion throughout mitosis. We used centromeric fluorescent CENP-A GFP+ foci, which can represent either an individual chromosome or a small cluster of chromosomes, as a way to track chromosome positions. Using FACS, a subpopulation of RPE1 cells with high GFP expression (enriched RPE1, eRPE1) was used to maximize the duration of live cell imaging against photobleaching. We examined centromeric movement/fluctuation in 4D. In particular, the degree of crossing three planes that defined by the centrosome-centrosome axis was quantified as shown (Fig. 2) as defined by our coordinate system, and the centrin1 protein in eRPE1 cells [17] (S2 Fig B,F,J).

**Fig. 2.**
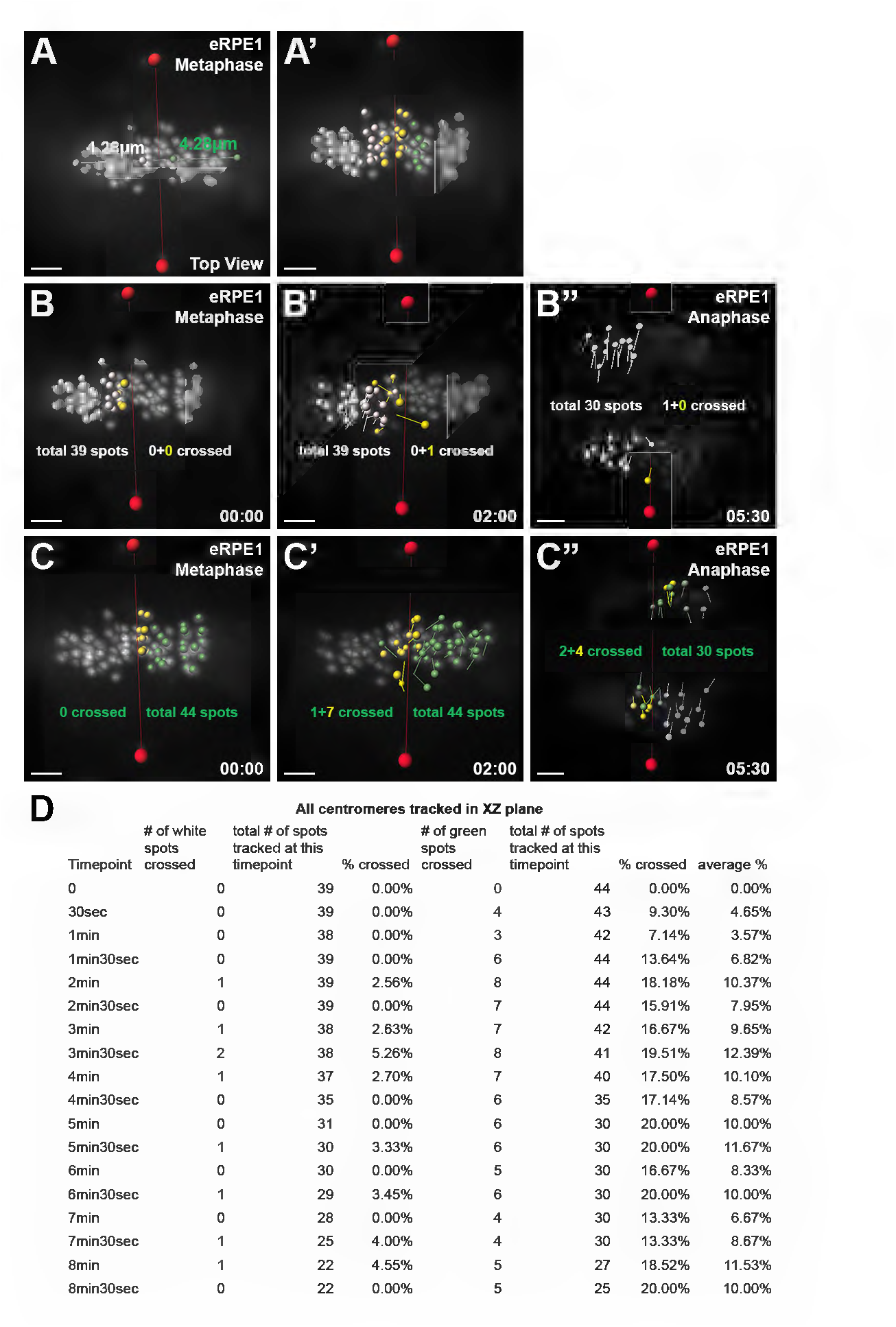
Centromere group identification and tracking of centromeres. Centromere group identification along the centrosome axis, or the XZ-plane (red line) of a metaphase enriched CENP-A/centrin1 GFP Retinal Pigment Epithelial (eRPE1) cell. **(A)** For easier visualization for centromere crossing, the long axis of the metaphase chromosome mass was divided into five equal parts. **(A’)** Centromeres positioned on either side of the XZ-plane, or centrosome axis (red line), were assigned the white/green group, while centromeres positioned in the middle ⅕ region (yellow) is used for tracking analysis. **(B-B’’)** Steps for tracking of centromere crossovers (%). Select frames of GFP+ centromeres, or spots, of eRPE1 cells within the white group (yellow) crossing over the XZ-plane (red line). The color and number of yellow spots crossing the XZ-plane at each time point was recorded, and divided by the total number of spots (%). Note: the total number of spots tracked decreased over time due to photobleaching. **(C-C’’)** As in (B-B’’) but of the centromeres within the green group (yellow). **(D)** Table of tracked centromeres that crossed the plane at each time point. GFP+ centromeres, or spots, for the white/green groups (yellow) were calculated at each time point, and averaged (%). Scale bar: 3 µm.

Tracking of centromere trajectories with CENP-A positive foci from metaphase to anaphase were performed with 30 sec acquisition intervals to minimize photobleaching (Fig 2 A,A’). Centromere group identification at either side of the XZ-plane of the centrosome axis of a metaphase eRPE1 cell was determined (Fig 2 A,A’). A centromere group was differentially tagged in white or green spots for GFP+ foci using an imaging application, IMARIS, as the two haploid chromosome sets are segregated [17]. Quantification of centromere crossing the XZ-plane of the centrosome axis was determined (Fig 2 B-D). For each timepoint, GFP+ centromeres, or spots, within the white/green groups that crossed over the XZ-plane were recorded, divided by the total number of spots, and averaged (%) (Fig 2 D).

The assigned groups of centromeres (white, or green, spots based on their initial positions) were tracked until anaphase and analyzed using defined parameters (Fig 3 A-B’). The centromere trajectories showed little, to no, mixing between the two sub-groups during this window of the cell cycle (Fig 3 C). While a minor overlap between the two groups was detectable, the composite tracks of the two groups are mostly separated from metaphase to anaphase (Fig 3 C). A result consistent with our previous report of continuous segregation of the two chromosome groups to either side of the XZ-plane of the centrosome axis [17].

**Fig 3.**
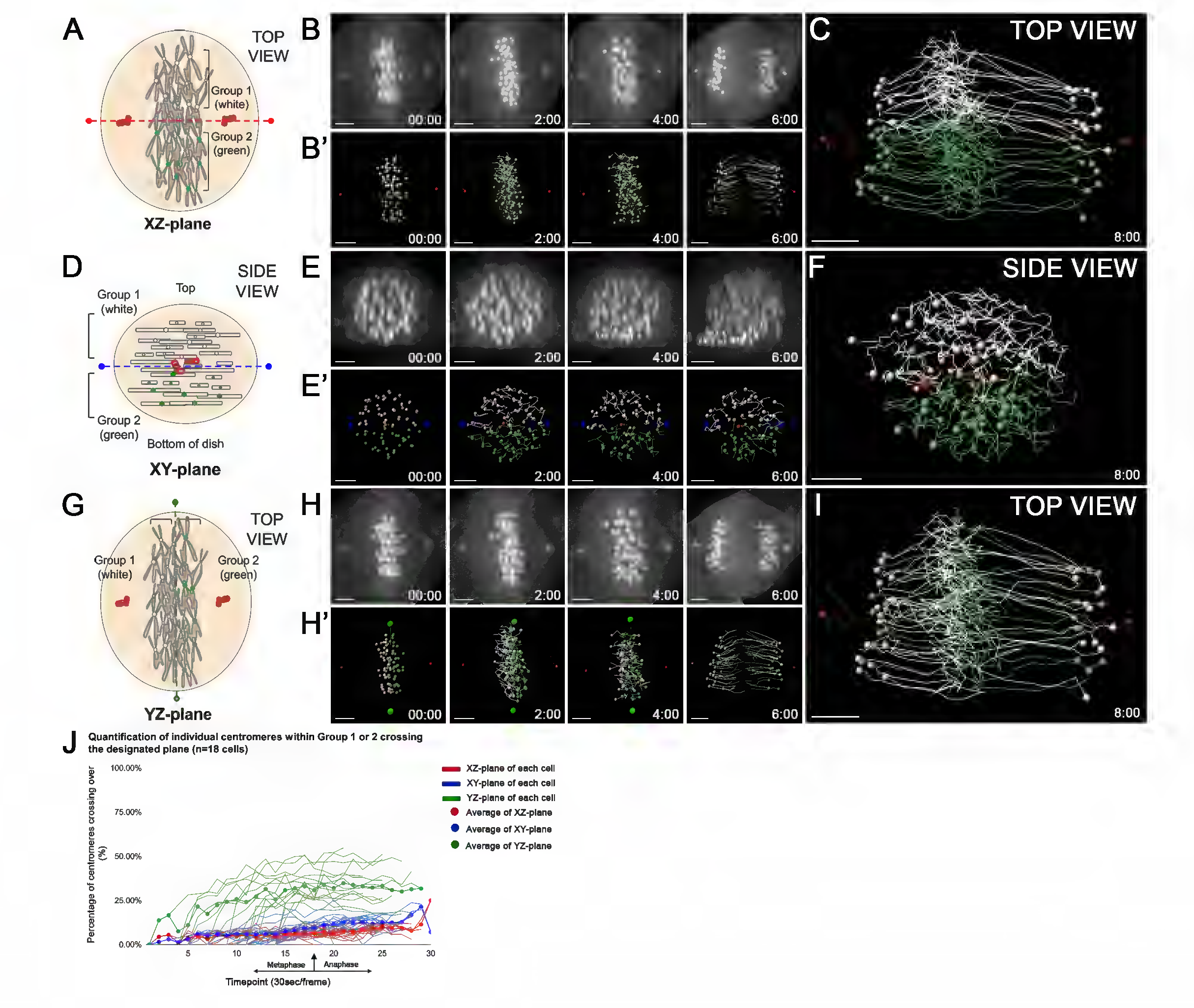
Ipsilateral, and apical/basal axis-based restriction of centromere movements along the centrosome axis from metaphase to anaphase. **(A)** Schematic of a human eRPE1 cell at metaphase. The XZ-plane (dashed red line) is defined as the plane connecting the centrosomes (red) and is perpendicular to the cell culture dish. Centromeres are assigned into two groups (white/green) based on their positions on either side of the XZ-plane. **(B)** Select frames of a time-lapse movie of a human eRPE1 cell from metaphase to anaphase acquired at 30 sec intervals. **(B’)** Tracking of centromeres (white/green spots) and centrosomes (red spots) in (B) to the XZ-plane (red line). Note: Lines following spots show the position of the centromeres and centrosomes from the previous two time points. **(C)** Composite trajectories of the two centromere groups (white/green spots) and centrosomes (red spots) labeled at metaphase, and tracked (white/green/red lines) to anaphase in a human eRPE1 cell for 8 min. **(D-F)** As in (A-C) but a side view showing centromere groups along the XY-plane (dashed blue line), which is defined as the plane parallel to the bottom of the cell culture dish. **(G-I)** As in (A-C), but of centromere groups along the YZ-plane (dashed green line), which is defined as the plane perpendicular to the XZ-plane, and the bottom of the cell culture dish. **(J)** Quantitative plot of centromeres (%) that cross over the XZ-plane (red), XY-plane (blue), and YZ-plane (green) from metaphase to anaphase (n=18 cells). The arrow indicates anaphase onset for cell alignment. Scale bar: 3 µm.

To test potential crossing of centromeres in other 3D planes of the centrosome-centrosome axis, centromeres were grouped based on their positions on either side of the XY-plane, or the apical-basal axis, and tracked (Fig 3D and S2 C,G,K Fig). The height along the apical/basal axis of the metaphase chromosome mass was measured, and the centromere groups were assigned into white and green groups similarly, as for the XZ-plane (Fig 3D-E’). Again, the data showed no detectable crossing between the two groups for the XY-plane, similar to the data seen in the XZ-plane (Fig 3A-E’). Examination of the centromere positions in the two groups showed that the centromere movements were restricted within the apical, and basal halves of the cell (Fig 3D-E’). The composite trajectories displayed a clear separation between the two groups (Fig 3F).

The YZ-plane is perpendicular to both the XZ-, and XY-planes (Fig 3G and S2 D,H,L Fig). Unlike the lack of crossing of the XZ- and XY-planes, the composite trajectories of the two groups exhibited highly detectable crossing of the YZ-plane (Fig 3H-I). Chromosomes continuously oscillate along the centrosome axis at metaphase alignment, before segregation to the daughter cells [40]. The YZ-plane serves as a control for a plane that utilizes a chance attachment to microtubules from one spindle pole to another for chromosome alignment at metaphase [40]. The data suggest that centromere movements are ipsilateral, and apical-basal restricted, specifically along the corresponding XZ- and XY-planes, but not in the YZ-plane.

To quantify the above motion data, centromere trajectories within the two groups at various time points for the three planes were analyzed, and the average percentage of white and green centromeres that crossed the designated planes were plotted (n=18 cells) (Fig 3J, and Table S1). The quantification analysis revealed that centromeres display restricted movements along the XZ-, and XY-planes, with an average of 6.6% , and 8.2% centromere crossovers (Fig 3J). In contrast, the YZ-plane displayed an average of 26.0% centromere crossovers (Fig 3J). This result is consistent with our previous data [17], and others [41–42] showing restricted centromere movements when attached to the mitotic spindle. Taken together, the data support an ipsilateral, apical/basal axis-based restriction parallel to the centrosome axis from metaphase to anaphase. Centromere motion restriction along both the x- and y-axes, dispute our previously published interpretation, leading to the likely possibility that motion restriction is largely facilitated by the mitotic spindle, rather than a haploid set-based segregation from metaphase to anaphase. Thus, we extend our analyses to non-spindle stages, in particular pre- and post-spindle stages. We next investigate centromere positions following mitotic spindle disassembly to test whether chromosome fluctuations are impacted.

### Restricted oscillation of centromeres at telophase and G_1_ interphase after mitotic spindle disassembly

During metaphase/anaphase, the mitotic spindle has been reported to play a role in facilitating chromosome movements [41, 43]. Following mitotic spindle disassembly at telophase, there has been conflicting data for its impact on chromosome positions at G_1_ interphase [44–47]. To test whether the ipsilateral, apical/basal-based restriction of centromere movements is present in the absence of the mitotic spindle, we performed live cell imaging of the eRPE1 cells from telophase to G_1_ interphase. RPE1 cells spend up to 8 hours in G_1_ interphase [48]. Therefore, eRPE1 cells were imaged for a minimum of 8 hours. To minimize phototoxicity and photobleaching, time lapse movies were captured at 15-30 min frame intervals to allow for longer imaging duration.

Centromere signals of the daughter cells were grouped at telophase as previously described by their positions on either side of the three planes (Fig 3 A,D,G). The centrosome axis was difficult to be determined after telophase due to the separation of daughter cells, and disappearance of one centrosome during cytokinesis [1]. At telophase, the nuclear membranes reform, and the daughter cells will flatten at G**_1_** interphase [49]. Under our sub-confluent culture conditions, when there is an increased number of mitotic cells as compared to confluent culture, the majority of the daughter cells (n=12/14 cells) were motile (Fig 4A-F and S2, S3 Videos). Individual tracks of the two centromere groups showed that the tagged centromeres (white or green) did not mix with those of the other tagged group from telophase to G_1_ interphase indicating no detectable centromere crossing at the XZ-plane (Fig 4A-A’). Composite trajectories on either side of the XZ-plane showed collective, and restricted movement of the centromere groups (Fig 4B). To test whether the restricted centromere movements are also present in the other 3D planes, centromere tracking for initial groupings across the XY-, and YZ-planes were conducted (n=7 cells) (Fig 4C-F, and Table S2). Centromere positions of the two groups also showed no detectable crossing at the XY-, and YZ-planes (Fig 4C’,D, E’, F). This data set shows that centromere oscillations are restricted along all 3D planes from telophase to G_1_ interphase.

**Fig 4.**
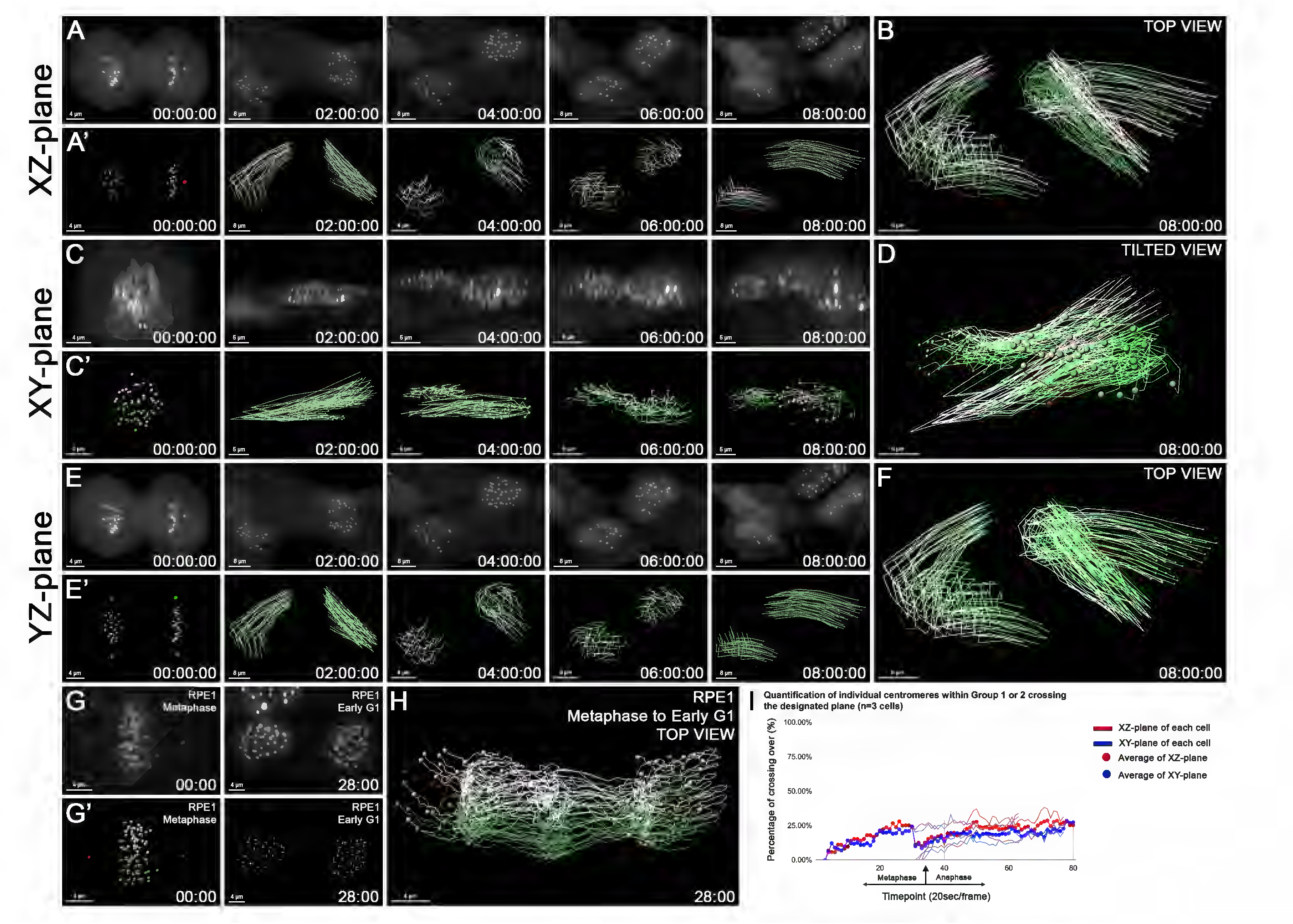
Restricted oscillation of centromeres at telophase and G_1_ interphase after mitotic spindle disassembly. **(A)** Select frames of a time-lapse movie of a human eRPE1 cell from late telophase to G_1_ interphase. **(A’)** Tracking of the centromeres (white/green spots) and centrosomes (red spots) in (A) to the XZ-plane. **(B)** Composite trajectories traveled by the two centromere groups (white/green spots) and centrosomes (red spots) labeled at telophase, and tracked (white/green/red lines) to G_1_ interphase in a human eRPE1 cell. **(C-D)** As in (A-B) but a side view (C), and tilted view (D) of identified centromere groups along the XY-plane. **(E-F)** As in (A-B) but of centromere groups along the YZ-plane. **(G)** An initial and final frame of a time-lapse movie of a human eRPE1 cell from metaphase, to early G_1_, with identified centromeres along the XZ-plane. **(G’)** Tracking of the centromeres (white/green spots) and centrosomes (red spots) in (G) labeled to the XZ-plane. **(H)** Composite trajectories traveled by the centromeres and centrosomes in (G) from metaphase to early G_1_. **(I)** Quantitative plot of centromeres (%) that cross over the XZ-plane (red) and XY-plane (blue) from metaphase to early G_1_ interphase (n=3 cells). The arrow indicates anaphase onset for cell alignment. Scale bars: 4, 5, or 8 µm

To capture centromere movements under the control of an intact spindle at metaphase to its disassembly at telophase, we performed live cell imaging for 1-1.5 hours. Time lapse movies with tagged centromere groups at either side of the XZ-plane were acquired at higher temporal resolution of 30 sec time frames to capture the dynamic chromosome movements (Fig 4G-G’ and S4 Video). Tagged centromeres were grouped at either side of the XZ-plane, at an initial time point at metaphase, and tracked to early G_1_ interphase (Fig 4G’). Two colored centromere groups had little crossover events after spindle disassembly. This result is consistent with the data from the time lapse movies with at least an 8 hour duration from telophase/G_1_ acquired at 30 min time frames (Fig 4A-F). Little crossing of the composite trajectories with white/green tagged centromeres confirmed that the ipsilateral, apical/basal axis-based restriction of centromere movements are preserved from metaphase to early G_1_ interphase (Fig 4H).

Quantification of potential crossing of centromere trajectories within the two groups at each timepoint for the XZ- and XY-planes were analyzed (n=3 cells) (Fig 4I and Table S3). The white/green centromeres that crossed the XZ- and XY-planes displayed movements along both planes, with an average of 20.1% centromere crossovers at the XZ-plane, and 17.1% at the XY-plane (Fig 4I). The YZ-plane was not analyzed at stages following anaphase/telophase, because sister chromatids have separated into the two daughter cells. Thus, it was technically difficult to reestablish the YZ-plane. This data set suggests that the restricted centromere oscillations within the two chromosome sets are conserved from metaphase to G_1_ interphase, even after spindle disassembly.

### Ipsilateral-based restricted centromere movements at prophase prior to mitotic spindle assembly

The ipsilateral, apical/basal axis-based restriction of centromere movements from metaphase to G_1_ interphase prompted us to extend our analysis of spatial regulation to mitosis onset, prior to spindle assembly [40]. To examine centromere positions and movements prior to spindle formation, we conducted a retrograde live cell imaging analysis from prophase to early metaphase (Fig 5A and S5 Video). At prophase, chromosomes undergo condensation, and become structurally visible in the cell [50]. Prior to metaphase, it was difficult to establish a reproducible axial coordinate system as centrosomes undergo dramatic positional changes [37, 17]. At prophase, the two centrosomes did not have the same distance from the surface of the coverslip, and the spindle and optical axes are not orthogonal. Therefore, we first used the centrosomes to revise our coordinate system with the x-axis, the z-axis is perpendicular to the x-axis with the smallest angle to the optical path, and y-axis is perpendicular to both the x- and z-axes. Centromere groupings were then established at early metaphase by identifying the centrosome axis, as well as the XZ-, XY, and YZ-planes (Fig 5B-G). As prometaphase is shorter than metaphase, the imaging analysis was conducted at 15 sec intervals.

**Fig 5.**
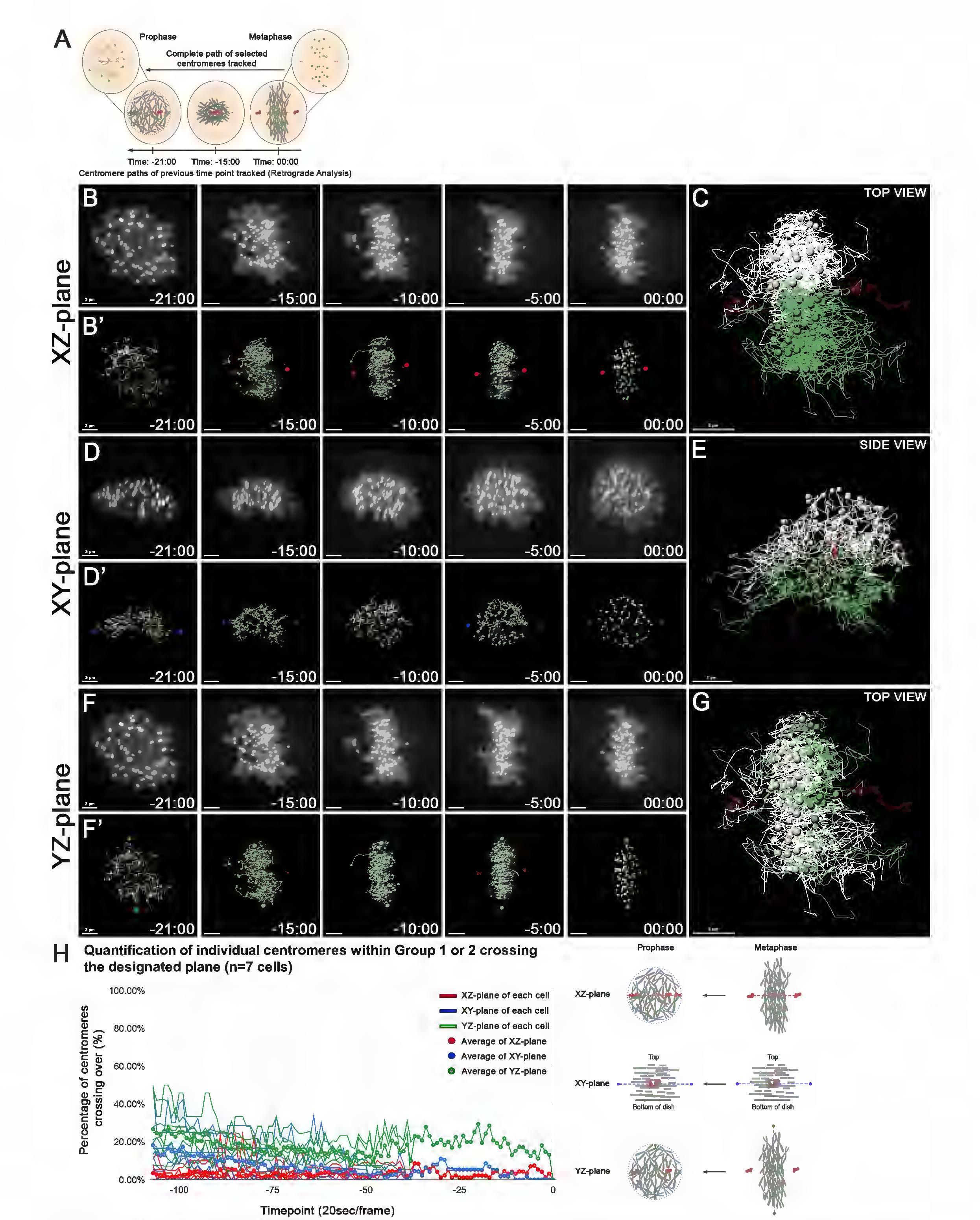
Ipsilateral-based restricted centromere movements at prophase prior to mitotic spindle assembly. **(A)** Schematic of retrograde tracking approach. Centrosomes (red) were identified at metaphase based on their positions outside of the chromosome mass, and tracked in reverse from metaphase to prophase. Centromeres were assigned into two groups (white/green spots) at metaphase, and tracked backwards. **(B)** Select frames of a time-lapse movie of a human eRPE1 cell from prophase to metaphase. **(B’)** Tracking of the centromeres (white/green spots) and centrosomes (red spots) in (B) to the XZ-plane (red line). Note: Lines following spots show the position of centromeres since the last two time points. **(C)** Composite trajectories of the two centromere groups (white/green spots) and centrosomes (red spots) labeled at metaphase, and retrograde tracked (white/green/red lines) to prophase in a human eRPE1 cell. **(D-E)** As in (B-C) but a side view of centromere groups along the XY-plane (blue line). **(F-G)** As in (B-C) but of centromeres along the YZ-plane (green line). **(H)** Quantitative plot of centromeres (%) that crossed over the XZ-plane (red), XY-plane (blue), and YZ-plane (green) from prophase to metaphase (n=7 cells). Schematic of centromere groups along the XZ-, XY-, and YZ-planes from prophase to metaphase. Scale bar: 3 µm.

Individual centrosomes were identified at early/mid-metaphase by their positions outside of the chromosome mass, and tracked back in reverse, or retrograde, to their positions at prophase (Fig 5A, S5 Fig) [51]. The centromeres were then tracked forward from prophase to early/mid metaphase, with their groupings established at early/mid-metaphase (Fig 5B’,D’). Examination of the centromere positions at each time point revealed that the centromere groups did not crossover the XZ-, and XY-planes, and the composite trajectories showed that the two groups remain segregated from prophase to metaphase (Fig 5B-E). Centromeres were found to crossover at the YZ-plane (Fig 5F-G). Retrograde analysis data of centromere crossing at the YZ-plane served as a control for axis-based centromere movements for little crossover events at the XZ- and XY-planes. Dynamic chromosome movements at prometaphase, such as chromosome poleward oscillations, within the two regions at either side of the XZ- or XY-planes, and little crossing at these two planes suggest a spatially regulated restriction for centromeric motion.

We quantified the centromere trajectories, and the average percentage of centromere crossovers for the three planes (n=7 cells) (Fig 5H and Table S4). An average of 3.3%, and 6.3% of centromere crossovers in the XZ-, and XY-planes, respectively, as compared to 17.9% in the YZ-plane, indicating that the centromere groups are segregated along the centrosome, and apical/basal axes during this mitotic window as well (Fig 5H). Centromere movement patterns from prophase to early/mid metaphase are similar to those observed at metaphase/anaphase. Taken together, these results show that individual centromere movements are ipsilateral, restricted along the XZ-, and XY-planes, prior to mitotic spindle assembly, and remain restricted after its disassembly. Centromere movements and fluctuations that cross the XZ-plane, defined by the centrosome-centrosome and z-axes, occur less frequently than other planes. The data support an ipsilateral axis-based centromere restriction of chromosome sets that persists from mitosis onset to G_1_ interphase, even without a major contribution of the mitotic spindle.

## Discussion

This work reveals an ipsilateral, apical/basal axis-based restriction of centromere movements from mitosis onset to G_1_ interphase in human cells. Our data demonstrate, for the first time, a clear discrete distribution of centromere components of satellite DNA and CENP-B protein with two peaks and a valley across the centrosome axis (Fig 1). The region of low staining of DNA satellite and CENP-B centromeric components may be the boundary between the two chromosome sets in primary human cells. We have previously defined a haploid chromosome set organization across the centrosome axis in multiple cell types [17]. Taken together, the axis-based restriction of centromere movements from mitosis onset to G_1_ interphase suggest the persistence of haploid sets throughout the cell cycle in human cells (Fig 6).

**Fig 6.**
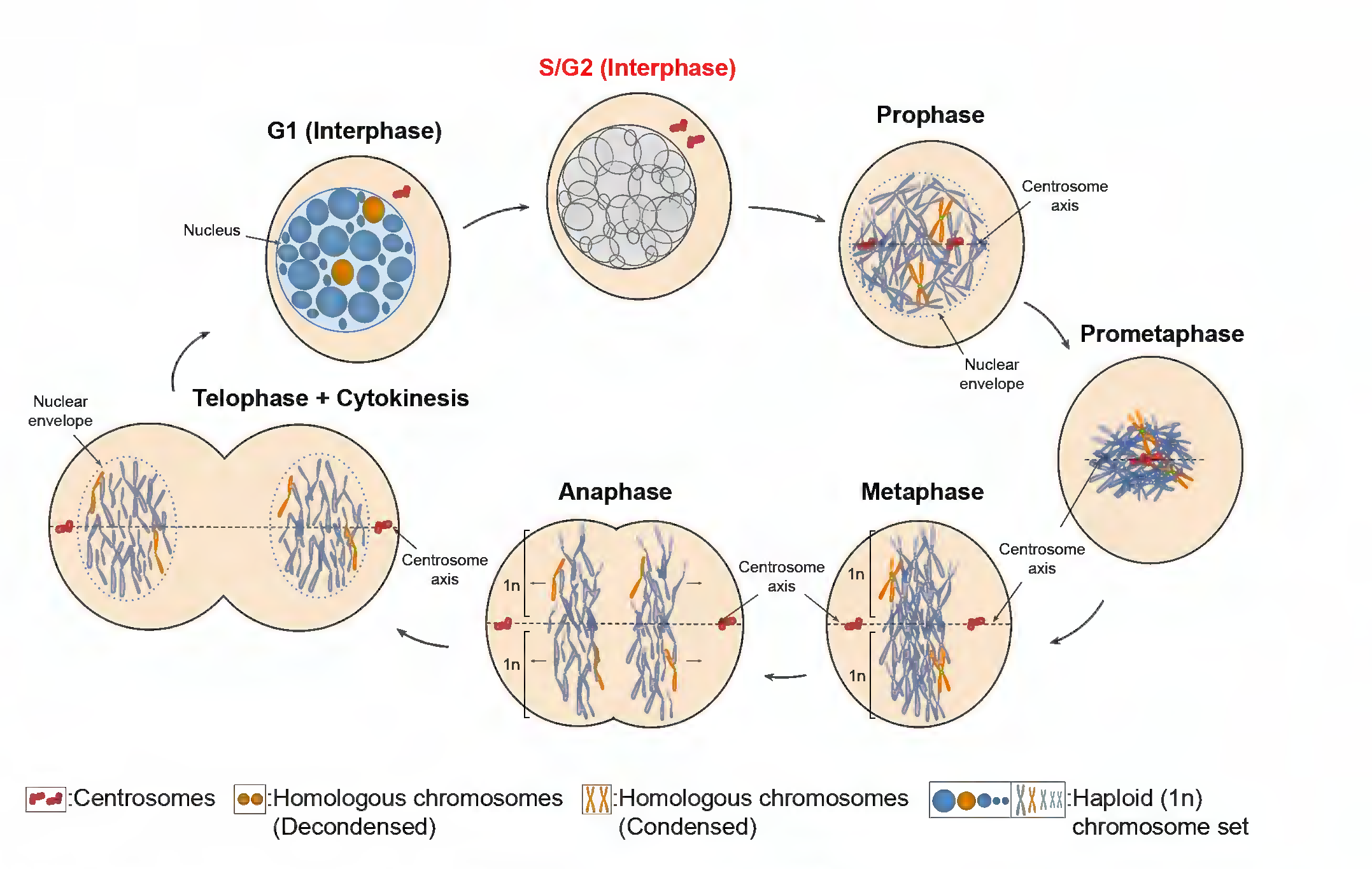
Chromosome movements throughout the cell cycle. Schematic of the XZ-plane, denoted by the centrosome axis, may be the boundary between two chromosome sets present throughout the cell cycle. Ipsilateral, apical/basal axis-based centromere movements are present from prophase to G_1_ interphase during mitotic spindle assembly, and after its disassembly. These data support a model that the haploid set organization may be present throughout mitosis onset to interphase.

The two discrete peaks of centromeric components found in the present study and the haploid set-based segregation of homologous chromosomes during cell division [2] lead to the question of how the spatial restriction of chromosome movement and positioning is regulated during the cell cycle. The current work is an initial step to address this obvious question by analyzing dynamics of centromere oscillation and movement. The descriptive work however leaves underlying mechanisms unsolved. It remains to be explored how this novel centromeric distribution pattern functions or is established as a boundary between chromosome sets. It would be plausible that inter-chromosomal linkages between the chromosomes of a haploid set may play a role for the unique centromeric pattern that may regulate haploid set organization. For example, centromeric linkages between individual chromosomes within each haploid set that may physically tether the chromosomes together. Pronuclear envelope proteins have been shown to contribute to parental genome separation in human and bovine embryos by forming a physical partition between parental genomes [52]. A similar molecular/cytoskeletal barrier could be acting to partition the haploid sets from each other in human cells. The region of low centromeric staining along the centrosome axis could then be a result of the bilateral segregation of the chromosome sets. However, there have been little to no studies describing such a molecular partition in human cells; therefore, closer investigation is needed.

Other mechanisms may be involved to regulate haploid set organization. Parental origin or epigenetic identity may also be implicated [17]. Previously, we have shown that a maternally derived translocation chromosome segregates within the same nuclear hemisphere as the X chromosome in male mouse fibroblasts [17]. This suggests that the individual haploid sets may be defined by parental origin. Separation of mammalian parental genomes at fertilization and early embryonic stages have been reported [53–54]. As such, a parental haploid set identity may be inherited, and transmitted throughout the development of the organism.

Individual chromosomes from metaphase to anaphase display ipsilateral, apical/basal axis-based restriction of movements along the centrosome axis (Fig 3). This result is consistent with our previous study [17] and others [37, 55]. However, our previous interpretation that the restricted centromere movements along the centrosome axis as a consequence of the haploid set organization needs to be revised. The current data suggest that the restricted centromere movements at this window of the cell cycle are likely a combination of mitotic spindle influence that contribute to a continued segregation of two chromosome sets during metaphase/anaphase.

Currently, there is a lack of consensus regarding chromosome behavior throughout the cell cycle. It is unknown whether individual chromosomes mix, or move collectively relative to each other. Using fluorescently labeled centromere, and centrosome CENP-A/centrin1 GFP Retinal Pigment Epithelial-1 cells, we showed a global maintenance of centromere position during the metaphase to anaphase progression along the centrosome axis [17]. Other studies utilized photobleaching experiments with labeled regions of interphase nuclei, and showed that chromosomes did not reorganize their positions throughout the cell cycle [44–45]. However, motion analysis of specific chromatin loci showed increased chromatin mobility from mitosis to G_1_ interphase during the cell cycle [46–47]. In these studies, different chromatin regions or loci were labeled and tracked, ranging from half or partial nuclei, or synthesized DNA of artificial LacO arrays, and native repeats. These may have contributed to different interpretations for individual chromosome behavior. In addition, cell type differences may also have contributed. Different cell lines including HeLa cells, human fibrosarcoma cells, CHO cells, and NRK epithelial cells were used for these studies, which may also result in variation in chromosome movements among cell lines.

Our data demonstrate that centromere fluctuations are restricted along the centrosome, and apical/basal axes present at prophase that persists until G_1_ interphase, even without major contribution of the mitotic spindle. Chromosome movements from the beginning to end of mitosis seem to be restricted suggesting the organization at mitosis onset may persist to G_1_ interphase. Individual chromosome positions have been suggested to vary in single cell populations [56]. Yet, to be studied is how a progenitor population has unique or distinct chromosome positions that can persist across multiple cell divisions and regulate a cell’s fate, behavior, and genetic fidelity. We cannot rule out the possibility that chromosome positions change during S/G_2_ interphase that may contribute to the varying positions of chromosomes.

In addition, nascent microtubules assemble between centrosomes prior to spindle formation during prometaphase [57–59] and may play a role in the chromosome axis-based restriction. Steric repulsion between chromosomes may also impact chromosome dynamics throughout the cell cycle and should be considered [60].

Our analysis utilizes the human RPE1 cell line to track individual or a small cluster of centromere positions, in contrast to tracking large subsets of chromosomes via photobleaching. We used centromeric, CENP-A foci, which is a specific and small domain for each chromosome, as a way to track chromosome positions throughout prophase to G_1_ interphase. To maximize the duration of live cell imaging, we enriched for a subpopulation of RPE1 cells with high GFP expression. The eRPE1 cell line showed homologous chromosomes 1 and X to be segregated along the centrosome axis at metaphase (S6 Fig). Chromosomes 4 and 13, however, showed a random distribution (S6 Fig), suggesting this eRPE1 cell line may not conserve the antipairing pattern for all chromosomes. We also acknowledge that the G_1_ phase nuclei are flat, and our tracking analysis along the XY plane may not be as precise as tracking in the other XZ- and YZ-planes as it does not account for the limitations of the axial optical resolution of our microscope.

While 4D high resolution live cell imaging allows us to map centromere trajectories throughout the cell cycle, it would be advantageous to label the homologous chromosomes. Our analysis tracks CENP-A GFP signals for centromeres over time, but does not provide identification of homologous chromosomes. Live imaging systems using fluorescent transcription activator-like effector (TALES), or Clustered regularly interspaced short palindromic repeats and CRISPR-associated protein 9 (CRISPR/Cas9) have been used to visualize centromeres/telomeres consisting of repetitive genomic sequences in mouse and human cell lines [61–62]. Although technically challenging due to the high condensation of chromosomes, chromatin inaccessibility, and photobleaching, labeling of homologous specific DNA loci will be necessary to definitively track chromosomes. It would be ideal to perform live cell imaging and tracking of centromere movements for multiple cell cycles to test cell viability, but such experiments are limited by photobleaching.

Loss or gain of function (LOF)/(GOF) assays for centromeric components to perturb the distribution pattern would be very difficult, if not impossible, at this time. We do not know yet how the discrete pattern is established and maintained throughout the cell cycle. In this study, we observe the distribution of α-satellite/cenpb sequences and the CENP-B protein. Centromeric α-satellite/cenpb sequences serve as the major structural DNA component of the centromeres [63–64]. As such, conducting perturbations to directly test the function of α-satellite/cenpb centromeric components for haploid set organization is not possible without impacting mitosis. Further study will be necessary to understand the relationship between the centromeric distribution pattern of two peaks and one valley, ipsilateral chromosome movements, and bilateral segregation of haploid sets along the centrosome, and apical/basal axes.

Our findings reveal new insights into chromosome organization and dynamics. We build upon the previous model of haploid chromosome set organization along a subcellular axis to describe an ipsilateral, apical/basal axis-based restriction of chromosome movements along the centrosome-centrosome axis in human cells. Implications of our study will contribute new knowledge of genome organization and shed light onto mechanisms that are implicated in human disease.

## Materials and methods

### Cell culture

Primary human umbilical vein endothelial cells (HUVECs, ATCC: PCS-100-013), and the human retinal pigment epithelial cell line (CENP-A/centrin1-GFP RPE1) were grown and cultured as previously described [17].

### Cell identification

Fluorescence-activated Cell Sorting (FACS) was performed for enrichment of a CENP-A/centrin1-GFP RPE1 cell population with high GFP expression (eRPE1). The eRPE1 cell line [37] is a heterogeneous population co-expressing CENP-A-GFP and centrin1-GFP.

Cells were trypsinized, and spun down to form a cell pellet, and resuspended in 1X PBS with 5% FBS. Cell counts were then performed (1.3x10^6^ cell/ml actual concentration) before transferring the cell suspension into a 5 ml round-bottom FACS tube with a 35 µm cell-strainer cap (Falcon, Cat#352235). Cell suspension was kept on ice until sorting. Sony SH800 FACS (UCSF) was used to measure the distribution range of GFP fluorescence. Cells were bulk sorted for the top 50-85% GFP fluorescent signal, and transferred into cell culture plates. Cells expressing the top 15% GFP fluorescence were excluded, as GFP overexpression is correlated with cellular defects, and death [65]. 70% of the sorted eRPE1 cells are diploid based on chromosome painting (data not shown).

For the eRPE1 cells, we found the duration of mitosis is ∼90 min as compared to 38-60 min in other RPE1 populations [66–67]. Distance between the centrosomes was used for staging of mitosis in the eRPE1 cell population with the average distance at prometaphase of 8.98 µm (n=7 cells) and early/mid-metaphase at 12.75 µm (n=8 cells).

### Fixed cell imaging

Mitotic cells were identified by brighter DAPI fluorescence intensity compared to surrounding interphase cells, DNA morphology, and centrosome position. Prometaphase cells were identified by the wheel-like rosette organization of the chromosomes as described [6, 37–38, 68]. Metaphase cells were identified by chromosome alignment at the equatorial plate, with a centrosome on either side. Fixed mitotic HUVECs were imaged with a confocal microscope (Leica TCS SPE, DM2500) using a 63x/1.3NA oil immersion objective with a digital zoom of 1.5x. The image data were acquired sequentially in a four-channel mode. Z-stacks were captured using a frame size of 1,024 x1,024 pixels, and processed with Leica Application Suite X software (Version: 3.5.2.18963). Confocal optical sections were reconstructed and visualized in Imaris software (Bitplane: 9.8.2). Imaris deconvolution algorithm (iterations: 10, pre-sharpening gain: 7.0) was applied, and the fluorescence level for all channels was thresholded by including 90% of each signal.

### Live cell imaging

eRPE1 cells were grown on 35-mm glass bottom µ-dishes (ibidi, Cat#50-305-807). Cells were imaged in culturing media; 37°C and 5% CO_2_ was maintained using a cage incubator and a stage top chamber (OkoLab). Time-lapse z-stack images were captured on an inverted Ti-E microscope (Nikon) equipped with a CSU-X1 spinning disk confocal (Yokogawa), motorized XY stage with Z piezo (ASI), 4 line laser launch (Vortan), Lambda 10-3 emission filter wheel (Sutter), quad-band dichroic ZET 405/488/561/640x (Chroma), with Plan Apo VC 100x/1.4NA and Plan Apo VC 60x/1.3NA oil objectives, and a Photometrics Prime95B sCMOS camera (Teledyne). eRPE1 cells were imaged using a 488-nm laser and ET525/50m emission filter (Chroma). z-stacks were acquired every 20-30 sec for 1 hr for mitotic cells (Figs 2, 3, 5) using the 100x/1.4NA objective, and every 15-30 min for up to 10 hours for interphase cells (Fig 4) using the 60x/1.3NA objective. Image processing was conducted using Imaris software.

### Immunolabeling and fluorescent in situ hybridization (ImmunoFISH)

HUVECs were grown on PTFE glass slides (Electron Microscopy Sciences, Cat# 63416-08) to 70-80% confluence for ImmunoFISH. Slides were washed with 1x DPBS (Gibco, Cat#14190250) twice before fixation with fresh 4% paraformaldehyde. After fixation, slides were washed 3X with chilled 1x PBS on ice, then underwent heat-induced antigen retrieval for 10 min in a sodium citrate buffer solution (10mM sodium citrate, 0.05% Tween-20, pH 6.0) in a steamer. Slides were then washed in a permeabilization buffer (0.25% Triton X-100 in 1x PBS) for 10 min followed by three 1x PBS washes at room temperature (RT). Slides were covered with a blocking buffer (10% goat serum, 0.1% Tween-20 in 1x PBS) for 30 min at RT. Slides were incubated with a primary antibody to γ-tubulin [1:1000, abcam: Anti-γ tubulin antibody (ab11317)] diluted in 0.1% Tween-20, 10% goat serum in 1x PBS. Slides were then covered in Parafilm, and incubated at 4°C in a humidified chamber overnight (O/N). The slides were washed 3X in 1x PBS. Slides were then incubated with a secondary antibody to goat anti-rabbit IgG [1:500, abcam: Goat Anti-Rabbit IgG H&L (Alexa Fluor® 647) (ab150079)] in 0.1% Tween-20, in 1% BSA and 1x PBS), covered in Parafilm, and incubated in the dark for 1 hr at RT. Slides were then washed twice with 1x PBS, and incubated with an EGS crosslinker solution (25% DMSO, 0.375% Tween-20, 25mM EGS in 1x PBS) for 10 min in the dark. Slides were washed twice with 1x PBS, and dehydrated via chilled ethanol series on ice (70%, 80%, 100%). DNA FISH probes to α-satellite (CENT-Cy3, pnabio, F3003) and cenpb (CENPB-Alexa488, pnabio, F3004) sequences were pre-warmed at 85°C for 10 min before use. Probes were diluted in a hybridization buffer [1:50, 60% ultrapure formamide (Fisher BioReagents, Cat# BP228-100), in 20mM Tris-HCl buffer], prior to slide incubation for 5 min at 85°C. Probes diluted in the hybridization buffer were added to slides, and incubated at 85°C for 10 min. Slides were transferred to a humidified chamber, and incubated in the dark for 2 hrs at RT. Coverslips were removed, and slides were washed in pre-warmed post-hybridization buffer (1% Tween-20, 10% 20x SSC in diH_2_O at 60°C) for 3X for 10 minutes at 60°C. Slides were counterstained with DAPI before mounting with ProLong Gold antifade (Invitrogen, Cat#P36930), and sealed with coverslips.

### Chromosome painting

Chromosome painting for HUVECs and eRPE1 cells was performed as previously described [17]. Whole chromosome paints for chromosome 1 in FITC, chromosomes 4 and 13 in Aqua, chromosomes 19 and X in Texas Red (Applied Spectral Imaging) were used. DNA was counterstained with SYTOX (Invitrogen, Cat# S11381), and mounted with ProLong Gold antifade (Invitrogen, Cat#P36930).

### 3D reconstruction and overlay, distance and angular orientation measurements of homologous chromosomes

Overlay generation of homologous chromosomes in multiple fixed metaphase HUVECs and eRPE1 was performed as previously described [17]. Calculations and plot generation for 2D positional analysis, and 3D distance and angular measurements between homologous chromosomes in fixed HUVECs were performed as previously described [17].

### Heatmap for DNA satellite sequences and CENP-B protein

Fixed HUVECs at metaphase stained for DNA satellite sequences (ɑ-satellite and cenpb) and the CENP-B protein were used to generate comprehensive heatmaps. 3D overlays were generated by overlaying individual cells along the centrosome, or x-axis, in Adobe Photoshop and the composite image was exported to ImageJ software to create a heatmap (ImageJ Interactive 3D Surface Plot plugin).

### 3D centromere and centrosome tracking

Centromere and centrosomes were identified as CENP-A/centrin1 GFP positive signals, and were defined as 0.5 µm diameter spots [69]. The automated spot object tracking algorithm in Imaris software was used to track centromere trajectories over time. For automated tracking analysis, individual centromere tracks were identified using the autoregression motion algorithm. The max distance between two centromere positions (initial and final), and its actual position for two consecutive time points was determined by averaging the distance traveled by random centromeres. The max distance is 1.5 µm for eRPE1 cells undergoing mitosis [37], and 8-10 µm for interphase cells due to longer time frame acquisition. Automated tracking was conducted for centromeres and centrosomes followed by manual correction.

For manual tracking, two consecutive time points were analyzed in 3D to determine each centromere position for the subsequent time point. Nearby centromeres, or cell edges were used as references. An example of manual tracking analysis of centromeres in interphase is shown (S7 Fig). Manual tracking analysis was conducted for telophase to G_1_ interphase analysis (Fig 4) as the distance traveled by the centromeres are ∼7-10µm per 30 min time frames.

For retrograde tracking analysis of centromeres, an automated algorithm with defined parameters was conducted first, followed by manual tracking as additional corrections and/or adjustments were required for accuracy. For example, when individual centromeric movements spanned 1.5 µm between two 15 sec timepoints. Centrosomes are in closer proximity to the centromeres at prophase with an average distance between the two centrosomes of 8.98 µM (n=7 cells). Therefore, manual identification at early/mid-metaphase was conducted with an average centrosome-centrosome distance of 12.74 µm (n=8 cells).

### Criteria of identification

Identification of CENP-A GFP positive foci in sequential time frames was determined by spot identification in Imaris software based on the following algorithm (Table S1-4). CENP-A GFP foci were ∼0.5 µm (S8 Fig) and were used for spot identification. CENP-A GFP foci represented either individual or a small cluster of centromeres. Due to photobleaching, some CENP-A-GFP signals were undetectable towards the end of live imaging. Thus, leading to less spots tracked for the complete duration of imaging. In addition, when two signals were very close in proximity to each other, only one spot was identified and tracked.

### Algorithm for: Prometaphase to early metaphase, and metaphase to anaphase

Enable Region Of Interest = false

Enable Region Growing = false

Enable Tracking = true

Enable Classify = true

Enable Region Growing = false

Enable Shortest Distance = true

[Source Channel]

Source Channel Index = 1

Estimated Diameter = 0.500 µm

Background Subtraction = true [Filter Spots]

“Quality” above automatic threshold [Tracking]

Algorithm Name = Autoregressive Motion

MaxDistance = 1.50 µm

MaxGapSize = 2

Fill Gap Enable = true [Filter Tracks]

“Track Duration” above 75.0 s [Classification]

[Event Setup]

### Algorithm for: Telophase to G1 interphase

Enable Region Of Interest = false

Enable Region Growing = false

Enable Tracking = true

Enable Classify = true

Enable Region Growing = false

Enable Shortest Distance = true [Source Channel]

Source Channel Index = 1

Estimated Diameter = 0.500 µm

Background Subtraction = true [Filter Spots]

“Quality” above 369 [Tracking]

Algorithm Name = Autoregressive Motion

MaxDistance = 8.00 µm

MaxGapSize = 3

Fill Gap Enable = true

[Filter Tracks]

“Track Duration” above 2000 s

[Classification]

[Event Setup]

### Quantification analysis of centromere crossing

The number of centromeres that crossed over the XZ-, XY-, and YZ-planes at each time point were divided by the total numbers tracked (%) (Figs 3J, 4I, and 5H). For example, if 3/35 green centromeres (8.57%) and 5/40 white centromeres (12.5%) crossed over the XZ-plane, the average would be 10.54% centromere crossovers for the time point. Live imaging videos of eRPE1 cells were aligned at anaphase onset (vertical arrow) for analysis.

### Video cell body outline tracing for live cell imaging

Manual outlining of cell bodies in each frame of centromere/centrosome tracking videos was applied using the PolyLineStroke function within the Fusion tab in DaVinci Resolve video editing software (Version 1.1.4 Build 9). Tracked cells of interest were outlined in green, yellow, or blue in each frame, while other cells in the frame were outlined in white. Following tracing of colored outlines, a custom Gamma Space under Image Source Gamma Space was applied to video [Gamma: 0, Linear Limit: 6.4, Linear Slope: 20, Remove Curve: checked] in addition to 3D Keyer filter, leaving only the colored outlines. Videos were edited with the original centromere/centrosome tracking video in DaVinci Resolve to produce side-by-side comparisons.

## Acknowledgements

We thank Dr. John Sedat for his helpful suggestions and feedback. We thank T.M. lab members (Jeanette Hyer, Rieko Asai, Zhiling Zhao) and L.L.H. laboratory members for their comments and suggestions, including Marcos Peech for assistance with CENP-B protein analysis. L.L.H dedicates this work to the memory of her daughter, Josephine Jereb. Live imaging data for this study were acquired at the Center for Advanced Light Microscopy-CVRI at UCSF on microscopes obtained using funding from the Research Evaluation and Allocation Committee, the Gross Fund, and the Heart Anonymous Fund. Part of this work is incorporated into a Master’s thesis (C.J.C., and G.Q.P). This work was supported in part by NSF RUI Award #2027746 (to L.L.H.), CSUPERB New Investigator Grant (to L.L.H.), SSU start up funds (to L.L.H.), NIH R01 HL148125 (to T.M.), NIH R01 HL153736 (to T.M.), and Smith Family Funds (to T.M.).

P.C., G.Q.P., and C.J.C. performed experiments; P.C., C.J.C., G.Q. P., and L.L.H. conducted analysis; P.C., C.J.C., G.Q.P., L.L.H., and T.M. wrote the manuscript.

## Supporting information

**S1 Video. Rotation of a metaphase HUVEC along the centrosome axis.** A 3D reconstructed HUVEC at metaphase stained for α-satellite (red), cenpb (green), and γ-tubulin (gray) rotating along the centrosome axis. Line denotes a region of low centromeric staining (α-satellite and cenpb) near the centrosome axis. Scale bar: 4 µm.

**S2 Video. Chromosomes move collectively from telophase to G_1_ interphase.** A 4D time lapse movie of a human eRPE1 cell from telophase to G_1_ interphase. Two groups of centromeres along the centrosome axis, or the XZ-plane, were labeled (white/green spots) at an initial time point at telophase, and tracked (white/green lines) until G_1_ interphase (Centrosomes: red). Cell body outlines of cells of interest (green/blue/yellow) and others (white) are shown. Scale bar: 3 µm.

**S3 Video. Chromosomes move collectively from telophase to G_1_ interphase.** A 4D time lapse movie of a different human eRPE1 cell from telophase to G_1_ interphase similar to [S2 Video]. Two groups of centromeres along the centrosome axis, or the XZ-plane, were labeled (white/green spots) at an initial time point at telophase, and tracked (white/green lines) until G_1_ interphase (Centrosomes: red). Cell body outlines of cells of interest (green/blue/yellow) and others (white) are shown. Scale bar: 10 µm.

**S4 Video. Individual centromeres show minimal mixing from metaphase to early G_1_ interphase.** A 4D time lapse movie of a human eRPE1 cell from metaphase to early G_1_ interphase. Two groups of centromeres along the centrosome axis, or the XZ-plane, were labeled (white/green spots) at an initial time point at metaphase, and tracked (white/green lines) until early G_1_ interphase. (Centrosomes: red). Cell body outlines of cells of interest (green/blue/yellow) and others (white) were shown. Scale bar: 5 µm.

**S5 Video. Chromosomes condense locally at prophase, and maintain their positions to early metaphase.** A 4D time lapse movie of a human eRPE1 cell undergoing mitosis from prophase to metaphase. Centrosomes (red) were identified at metaphase, and retroactively tracked to prophase. Two groups of centromeres along the centrosome axis were labeled (white/green spots) at metaphase, and retroactively tracked (white/green lines) to prophase. Scale bar: 2 µm.

---

**Fig S1.**
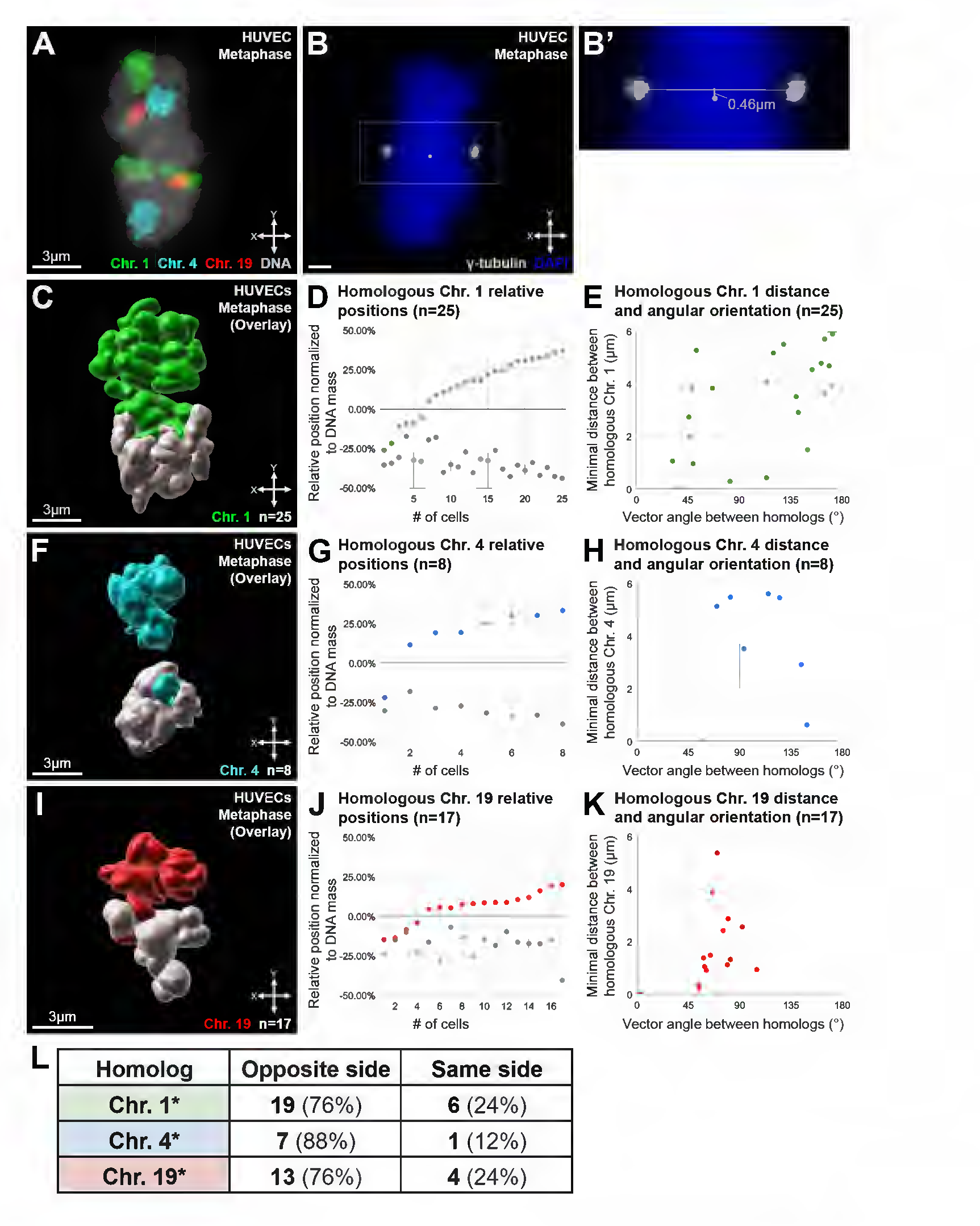
Antipairing of homologous chromosomes in Human umbilical vein endothelial cells (HUVECs). **(A)** A HUVEC at metaphase painted for chromosomes 1 (green), 4 (cyan), 19 (red), and DNA counterstain (SYTOX, gray). **(B)** A metaphase HUVEC stained for γ-tubulin antibody (gray), and DNA counterstain (DAPI, blue). Gray dot represents the center of mass of the DAPI stained chromosomal mass. **(B’)** Zoom in view of the region in (B). The distance between the centrosome axis (line connecting the centrosomes) and the DAPI center of mass on average is 0.5 μm. Thus, a 1μm width bounding box region of DAPI staining overlapping the DNA center of mass was determined as the “boundary zone.” **(C)** Top view of a 3D overlay of chromosome 1 (green/gray) distribution of multiple HUVECs at metaphase (n=25). Of a homologous pair, the individual chromosome 1 was assigned to be either green or white based on its proximity to the y-axis, when y=0 (green/white was most proximal/distal, respectively) and used to generate 3D overlay. **(D)** Relative positions for each pair of homologous chromosome 1 (green/gray circles) when mapped to an axial coordinate system (n=25). **(E)** Distance and angular orientation of homologous chromosome 1 pairs (n=25) with an average distance of 3.1 μm and angular orientation of 104.1°. **(F)** As in (C), but for chromosome 4 (cyan/gray) (n=8). **(G)** As in (D), but for chromosome 4 pairs (blue/gray circles) (n=8). **(H)** As in (E), but for chromosome 4 pairs (n=8) with an average distance of 3.6 μm and angular orientation of 104.4°. **(I)** As in (C), but for chromosome 19 (red/gray) (n=17). **(J)** As in (D), but for chromosome 19 pairs (red/gray circles) (n=17). **(K)** As in (E), but for chromosome 19 pairs (n=17) with an average distance of 1.7 μm and angular orientation of 64.5°. **(L)** Table for the quantification of homologous chromosome positions along the DNA center of mass, and XZ-plane. Note: Cells with a chromosome whose center of mass was positioned within the boundary zone were unable to be analyzed. Binomial probability of p=0.005* for chromosome 1, p=0.031* for chromosome 4, and p=0.018* for chromosome 19. Scale bar: 2 μm.

**Fig S2.**
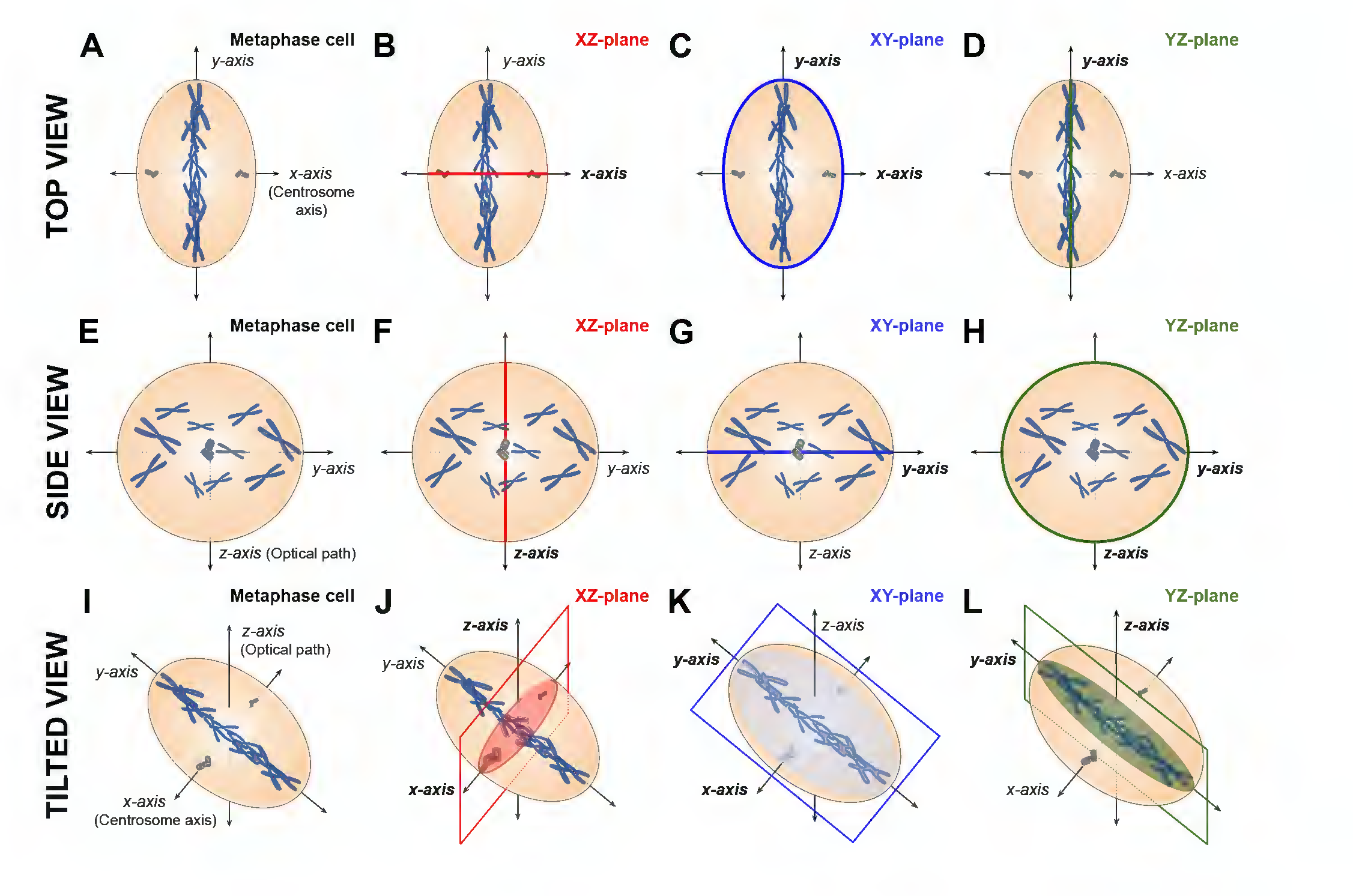
Subcellular coordinate axis system and planar divisions. **(A-D)** Top view, **(E-H)** side view, or **(I-L)** tilted view of schematic of a metaphase cell outlining the x-axis crossing the centrosomes, the z-axis following the optical path through the cell, and the y-axis perpendicular to both the x- and z-axes. **(B, F, J)** The XZ-plane (red outline) is defined as the plane connecting the centrosomes (gray) and is perpendicular to the cell culture dish. **(C, G, K)** As in (B, F, J), but showing the XY-plane (blue outline), which is defined as the plane parallel to the bottom of the cell culture dish. **(D, H, L)** As in (B, F, J), but showing the YZ-plane (green outline), which is defined as the plane perpendicular to the XZ-plane and the bottom of the cell culture dish.

**Fig. S3.**
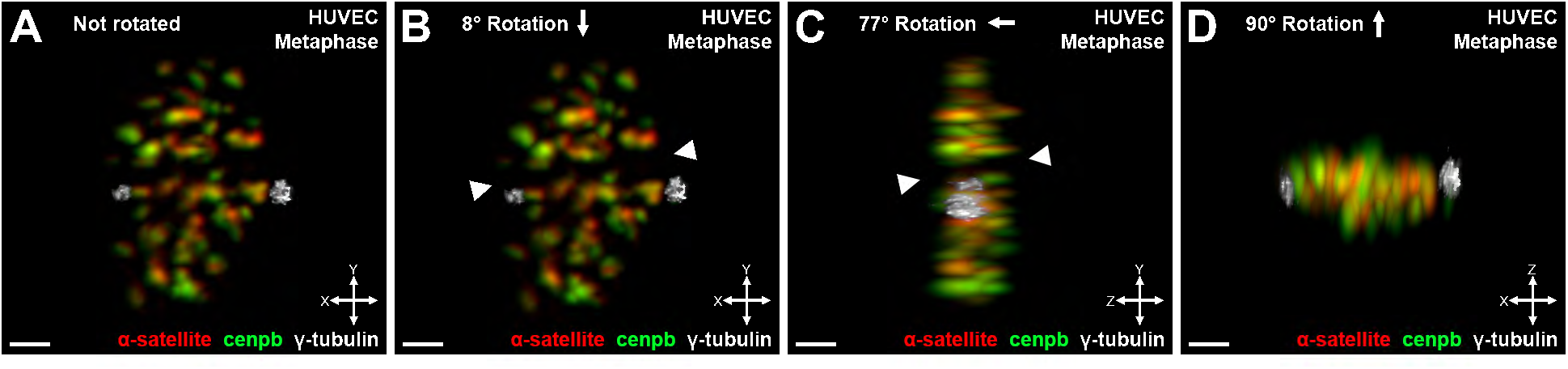
3D rotation analysis of metaphase HUVECs reveal a region of low DNA α-satellite and cenpb box staining along the centrosome axis. (A) Similar to [Figure 1B], but of a different metaphase HUVEC. (B) The HUVEC in (A) rotated 8° along the x-axis. (C) As in (B), but rotated 77° along the y-axis. (D) As in (B), but rotated 90° along the x-axis. Note: Rotation from multiple 3D perspectives demonstrates a region of low centromere staining for α-satellite and cenpb box (white arrowheads) along the centrosome axis. Scale bar: 2 µm.

**Fig S4.**
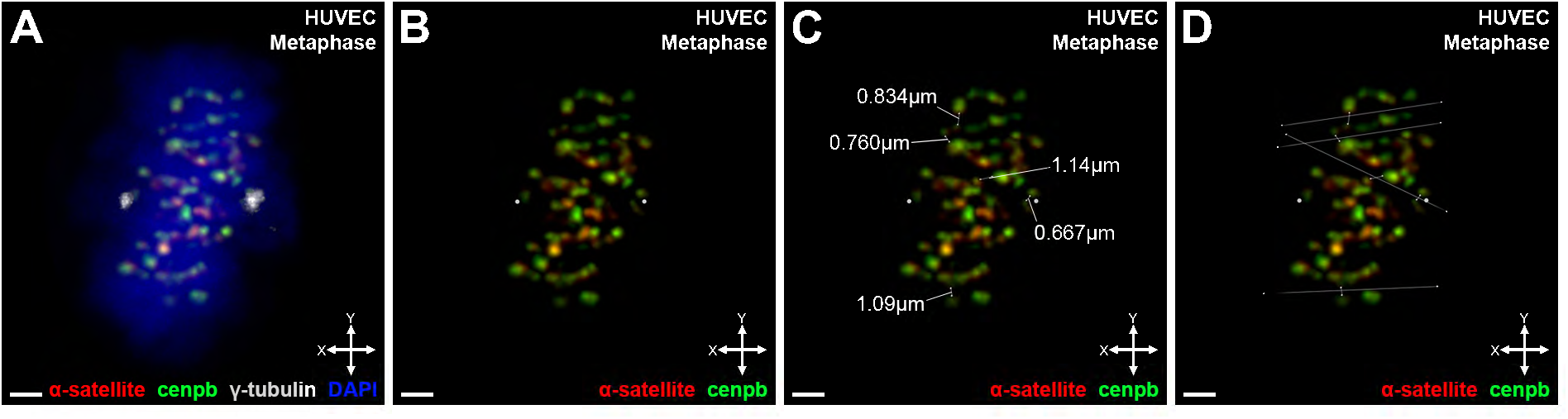
Characterization of multiple regions of low DNA satellite staining in a metaphase eRPE1 cell. **(A)** Same as [Figure 1B], but for another metaphase HUVEC. **(B)** As in (A) but without the DNA counterstain (DAPI). **(C-D)** Multiple regions of low staining of centromeric staining (α-satellite and cenpb) can be identified throughout the metaphase chromosome mass. Individual cells may exhibit multiple regions of low DNA satellite staining. Scale bar: 2 μm.

**Fig S5.**
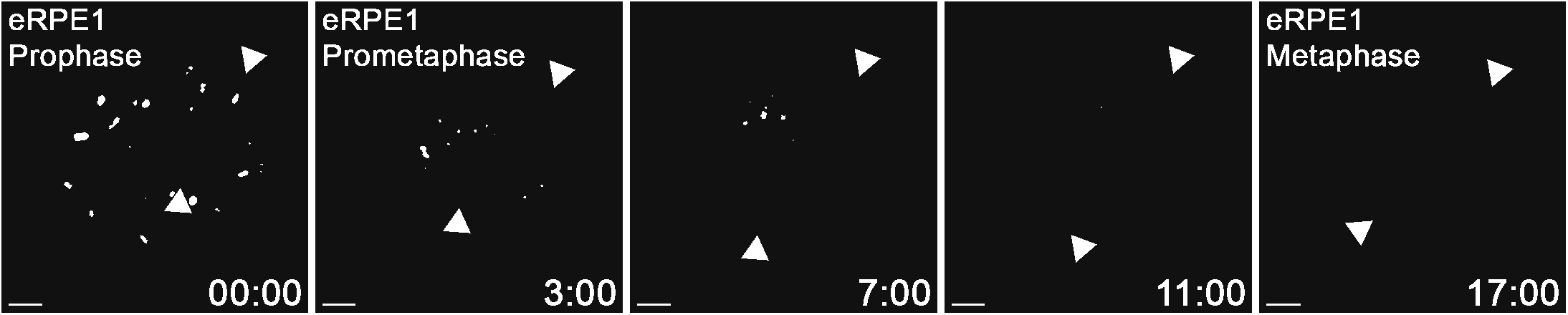
Manual retrograde tracking analysis of centrosomes from prophase to metaphase in an eRPE1 cell. As in [Fig 5], the centrosomes (arrowheads) were identified at metaphase by their location outside of the chromosome mass, then manually retrograde tracked to prophase. Scale bar: 2 µm.

**Fig S6.**
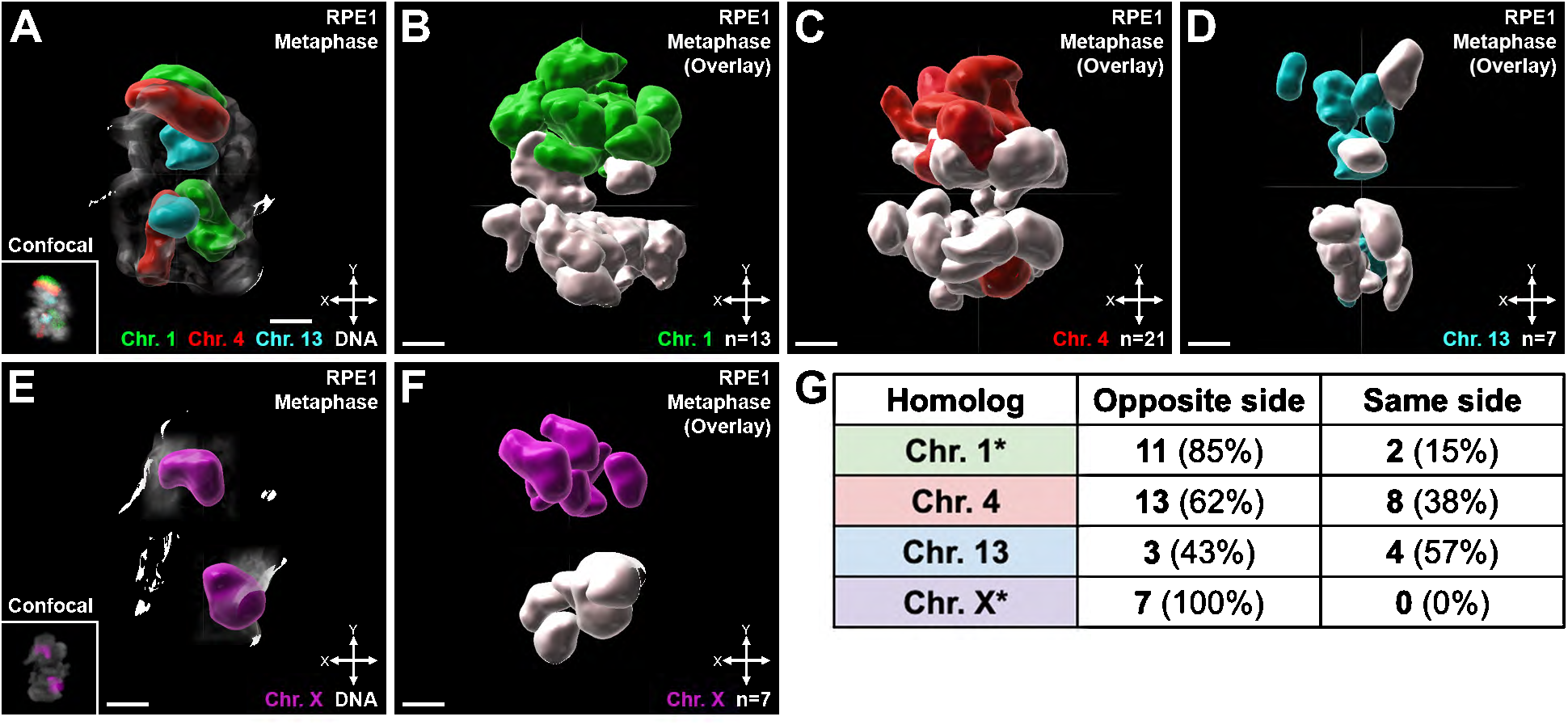
Chromosome analysis of eRPE1 cells. **(A)** A 3D reconstruction of a metaphase eRPE1 cell showing homologous chromosomes 1 (green), 4 (red), 13 (cyan), and DNA counterstain (SYTOX, gray). Inset: Stacked confocal optical sections of cell. **(B)** Top view of a 3D overlay of chromosome 1 (green/gray) distribution of multiple eRPE1 cells at metaphase (n=13). **(C)** As in (B), but for chromosome 4 (red/gray) (n=21). **(D)** As in (B), but for chromosome 13 (cyan/gray) (n=7). **(E-F)** As in (A-B), but for homologous chromosomes X in purple (n=7). **(G)** Table for the quantification of homologous chromosome positions along the DNA center of mass, and XZ-plane. Note: Similar to S1 Fig, cells with a chromosome whose center of mass was positioned within the boundary zone were not interpretable. Binomial probability of p=0.01* and p=0.008** for chromosome 1 and X. Scale bars: 2 µm.

**Fig S7.**
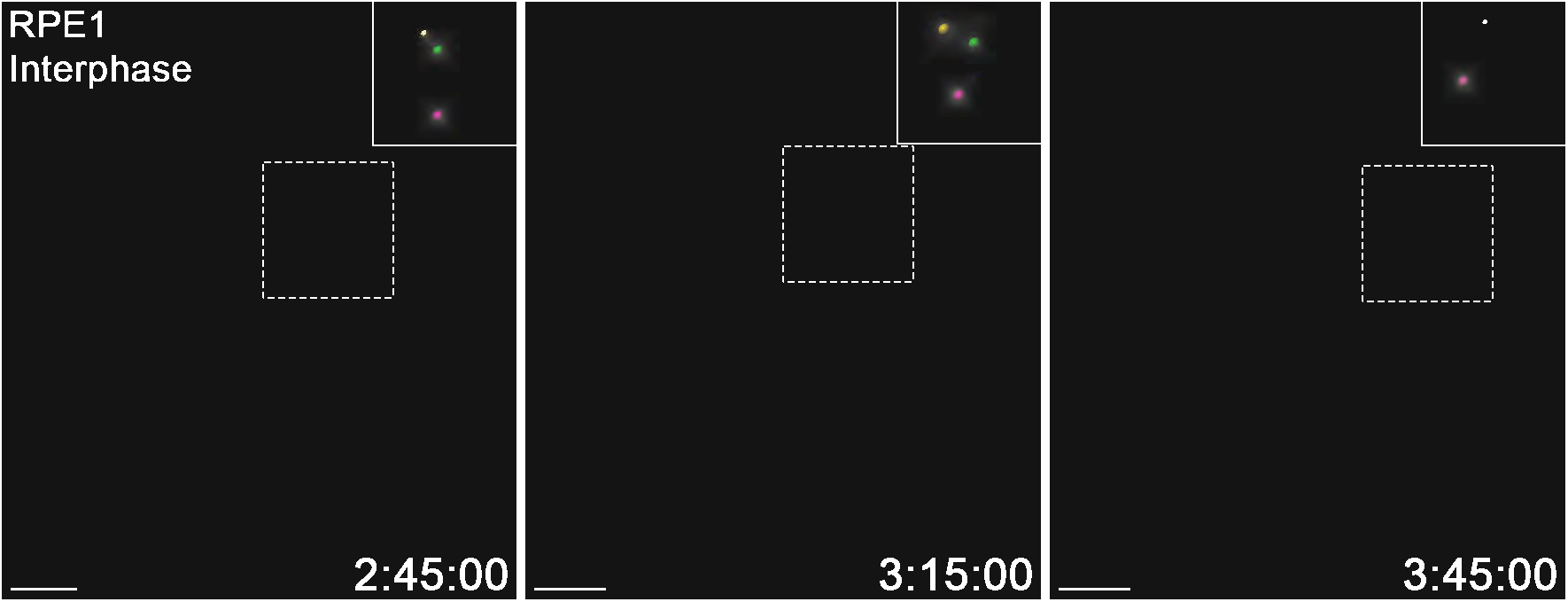
Manual tracking of interphase centromeres. Select frames of a time lapse video of a human eRPE1 cell at G_1_ interphase. Manual tracking of centromeres at interphase was performed by identifying individual centromeres (4-5 spots), and tracked over time. 4 different spots are shown as examples. Scale bar: 2 µm.

**Fig S8.**
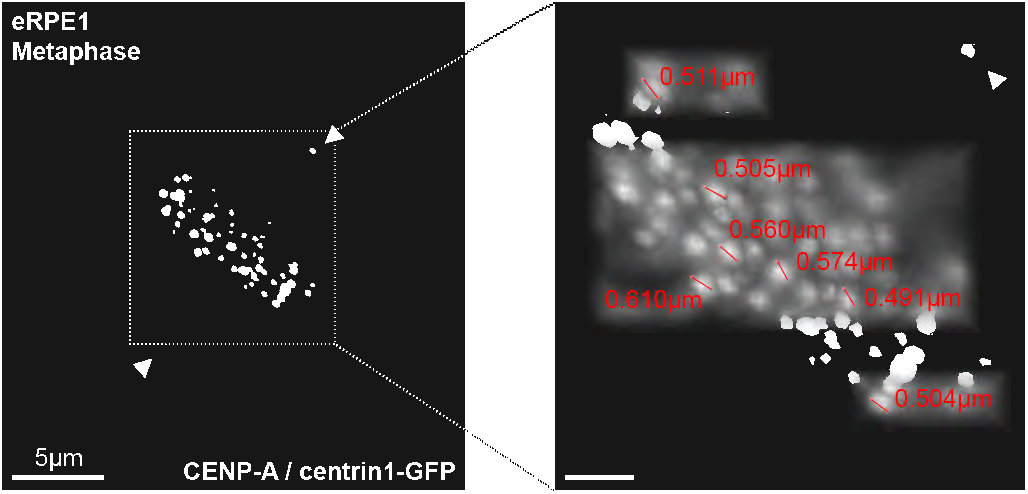
Quantification of CENPA-GFP foci of eRPE1 cells. A live metaphase eRPE1 cell showing individual measurements for multiple CENPA-GFP positive foci. On average, the CENPA GFP foci was 0.5 µm. Scale bar: 5 µm.

**Table S1.** Number of centromeres tracked for each eRPE1 cell used for live imaging analysis. Tables showing the total number of eRPE1 cells that were live imaged with the average number of spots, timepoints, and total duration during metaphase to anaphase.

| Stage: Metaphase to Anaphase (30sec intervals) |  |  |  |  |
| --- | --- | --- | --- | --- |
| Cell# | Average # of spots tracked per timepoint | Total timepoints | Total length tracked (sec) | Total length tracked (min) |
| 1 | 62 | 19 | 570 | 9.5 |
| 2 | 58 | 24 | 720 | 12 |
| 3 | 33 | 8 | 240 | 4 |
| 4 | 53 | 9 | 270 | 4.5 |
| 5 | 75 | 10 | 300 | 5 |
| 6 | 87 | 20 | 600 | 10 |
| 7 | 81 | 12 | 360 | 6 |
| 8 | 87 | 17 | 510 | 8.5 |
| 9 | 76 | 15 | 450 | 7.5 |
| 10 | 88 | 29 | 870 | 14.5 |
| 11 | 89 | 15 | 450 | 7.5 |
| 12 | 84 | 15 | 450 | 7.5 |
| 13 | 90 | 17 | 510 | 8.5 |
| 14 | 83 | 22 | 660 | 11 |
| 15 | 77 | 22 | 660 | 11 |
| 16 | 69 | 18 | 540 | 9 |
| 17 | 60 | 15 | 450 | 7.5 |
| 18 | 68 | 20 | 600 | 10 |
| Average | 73 | 17 | 512 | 9 |

**Table S2.**
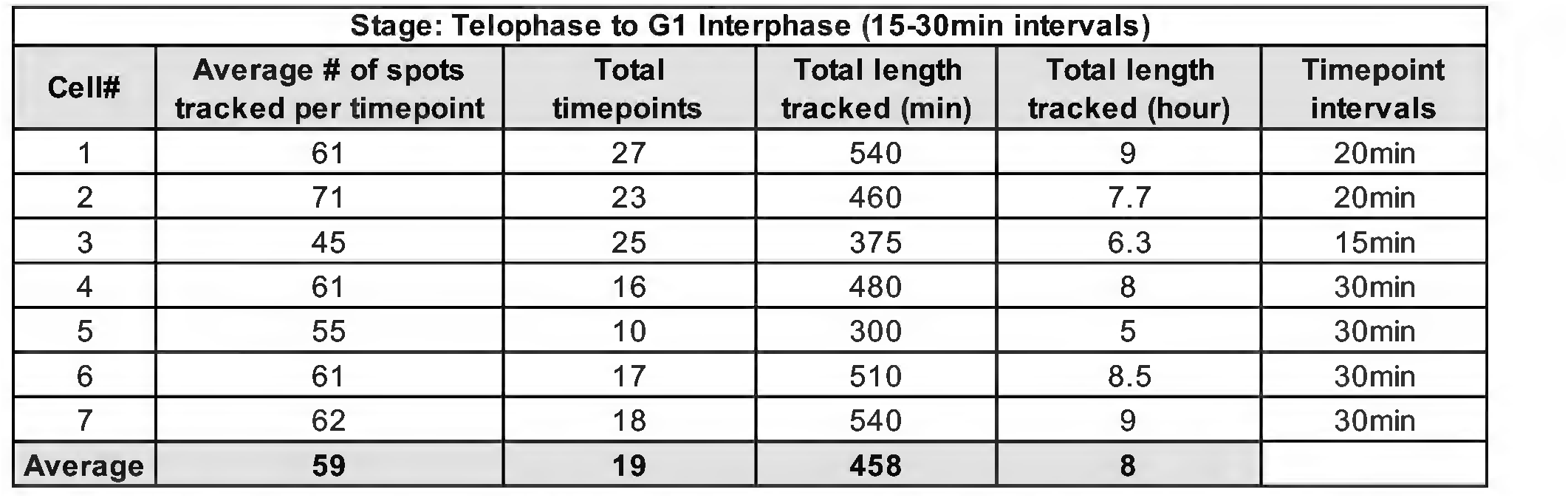
Number of centromeres tracked for each eRPE1 cell used for live imaging analysis. Tables showing the total number of eRPE1 cells that were live imaged with the average number of spots, timepoints, and total duration during telophase to G_1_ interphase.

**Table S3.** Number of centromeres tracked for each eRPE1 cell used for live imaging analysis. Tables showing the total number of eRPE1 cells that were live imaged with the average number of spots, timepoints, and total duration during metaphase to G_1_ interphase

| Stage: Metaphase to G1 Interphase (20-30sec intervals) |  |  |  |  |  |
| --- | --- | --- | --- | --- | --- |
| Cell # | Average # of spots tracked per timepoint | Total timepoints | Total length tracked (sec) | Total length tracked (min) | Timepoint intervals |
| 1 | 59 | 103 | 3090 | 51.5 | 30sec |
| 2 | 87 | 50 | 1300 | 21.7 | 26sec |
| 3 | 71 | 63 | 1260 | 21 | 20sec |
| Average | 72 | 72 | 1883 | 31 |  |

**Table S4.**
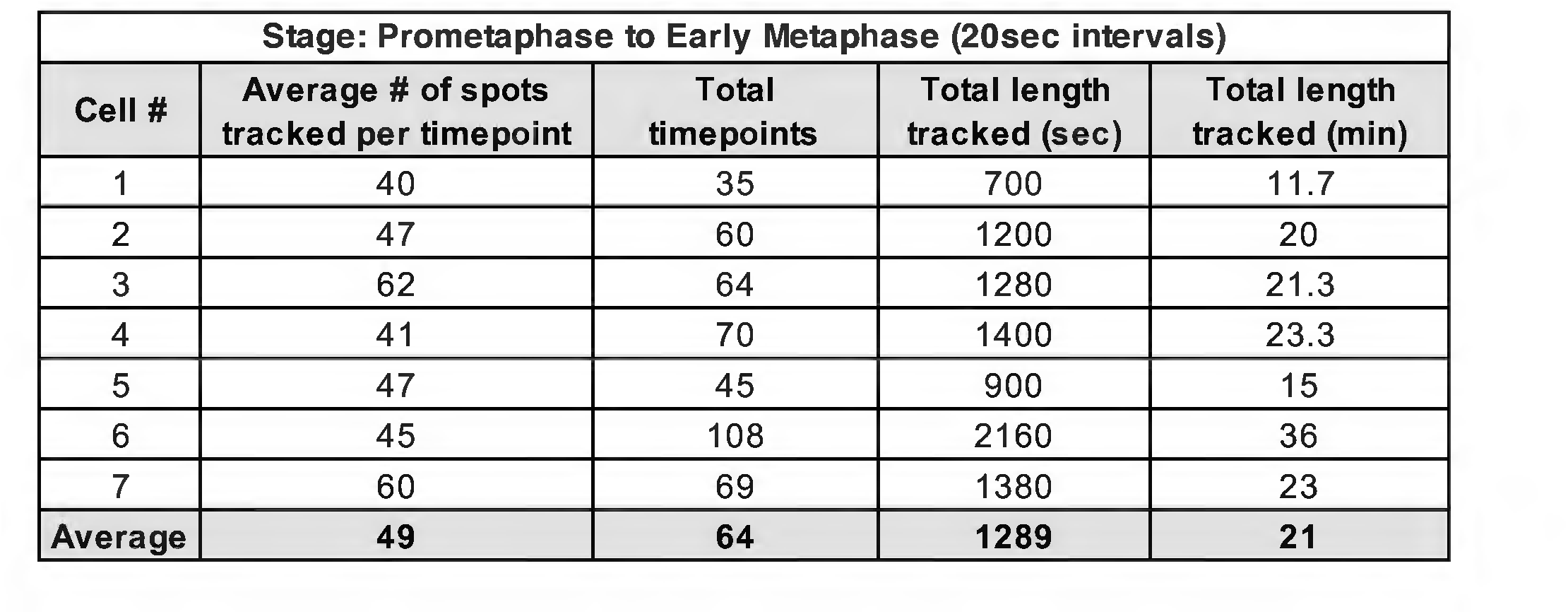
Number of centromeres tracked for each eRPE1 cell used for live imaging analysis. Tables showing the total number of eRPE1 cells that were live imaged with the average number of spots, timepoints, and total duration during prophase to early metaphase.

| Stage: Prometaphase to Early Metaphase (20sec intervals) |  |  |  |  |
| --- | --- | --- | --- | --- |
| Cell # | Average # of spots tracked per timepoint | Total timepoints | Total length tracked (sec) | Total length tracked (min) |
| 1 | 40 | 35 | 700 | 11.7 |
| 2 | 47 | 60 | 1200 | 20 |
| 3 | 62 | 64 | 1280 | 21.3 |
| 4 | 41 | 70 | 1400 | 23.3 |
| 5 | 47 | 45 | 900 | 15 |
| 6 | 45 | 108 | 2160 | 36 |
| 7 | 60 | 69 | 1380 | 23 |
| Average | 49 | 64 | 1289 | 21 |

